# AutoCoEv – a high-throughput *in silico* pipeline for predicting inter-protein co-evolution

**DOI:** 10.1101/2020.09.29.315374

**Authors:** Petar B. Petrov, Luqman O. Awoniyi, Vid Šuštar, M. Özge Balcı, Pieta K. Mattila

**Affiliations:** Institute of Biomedicine and MediCity Research Laboratories, University of Turku, Turku, Finland; Turku Bioscience, University of Turku and Åbo Akademi University, Turku, Finland

**Author notes:** Correspondence (PBP); (PKM).

**Keywords:** co-evolution, correlation, protein, interaction, automation, pipeline

## Abstract

Protein-protein communications govern cellular processes via complex regulatory networks, that are still far from being understood. Thus, identifying novel interactions between proteins can significantly facilitate our comprehension of the mechanistic principles of protein functions. Co-evolution between proteins is a sign of functional communication and, as such, provides a powerful approach to search for novel direct or indirect molecular partners. However, evolutionary analysis of large arrays of proteins, in silico, is a highly time-consuming effort, which has limited the usage of this method to protein pairs or small protein groups. Here, we developed AutoCoEv, a user-friendly computational pipeline for the search of co-evolution between a large number of proteins. By driving 15 individual programs, culminating in CAPS2 as the software for detecting co-evolution, AutoCoEv achieves seamless automation and parallelization of the workflow. Importantly, we provide a patch to CAPS2 source code to strengthen its statistical output, allowing for multiple comparisons correction and enhanced analysis of the results. We apply the pipeline to inspect co-evolution among 324 proteins identified to locate at the vicinity of the lipid rafts of B lymphocytes. We successfully detected multiple strong coevolutionary relations between the proteins, predicting many novel partners and previously unidentified clusters of functionally related molecules. We conclude that AutoCoEv, available at https://github.com/mattilalab/autocoev, can be used to predict functional interactions from large datasets in a time and cost-efficient manner.

## 1. Introduction

The biological function of proteins is carried out through association and communication with various molecules, the majority of which are other proteins. Thus, screening for novel interactions, either direct or indirect, is of high importance for deciphering the complexity of protein networks. It has been shown that relations between proteins can be extrapolated from the evolutionary history of their genes, via in silico analysis of co-evolution [1,2].

The evolution of proteins is influenced by structural and functional constraints between amino acids, enforcing their adaptation in a concerted manner. Detecting intra- or inter-molecular co-evolution is regarded as a sign of functional co-dependence between residues within the same protein, or between sites belonging to different partners, respectively [3]. Various computational approaches for prediction have been described, among which are BIS2 [4], ContactMap [5], DCA [6], Evcouplings [7], GREMLIN [8], MISTIC [9], PKSpop [10] and CAPS2 [11]. Notably, a comprehensive large-scale study was shown recently by Cong et al for bacterial proteome [12]. However, many of the searches for inter-protein co-evolution have been confined to a relatively small number of partners, where an existing correlation has been initially anticipated [13–17]. If applied to large datasets, such computational approaches would demand a high degree of automation, an issue that we successfully address in this work.

Here, we developed an automated computational pipeline called AutoCoEv, for the large-scale screening for protein interactions, that is user-friendly and ready to use by the broader public. In the center of the workflow is CAPS2 (Coevolution Analysis using Protein Sequences 2) software, that compares the evolutionary rates between sites in the form of their correlated variance [11]. By driving 15 programs, AutoCoEv achieves a high level of automation and flexibility, as well as, processes parallelization, enabling the analysis of hundreds of proteins on a regular computer. We demonstrate the performance of the pipeline by analyzing 324 lymphocyte lipid raft resident proteins [18], identified in a proximity biotinylation screen, for their potential functional relation.

## 2. Implementation

The preparation pipeline for most coevolutionary analyses has a relatively simple concept. Typically, for each protein of interest, a multiple sequence alignment (MSA) is produced from its orthologues in different species, optionally combined with a phylogenetic tree. However, this process requires the correct identification of orthologues, their sequences retrieval and high-quality alignment, all tied together by various filtrations and file format conversions. Automating these steps also necessitates a robust quality-check during and after the process, all of which can present significant challenges if done manually. While developing AutoCoEv we paid significant attention to incorporate ways to evaluate the quality of the key preparatory steps and, finally, to evaluate the results for their robustness.

### 2.1 Command line interface, configuration and input

AutoCoEv is written in BASH and offers a simple menu-driven command line interface (CLI), in which the individual steps are enumerated (Figure 1a). Options for the programs that AutoCoEv drives, as well as filtering parameters, are configured in a single file (settings.conf), described in detail in the manual distributed with the script. Once configuration has been set, simply going through the steps consecutively will conduct the work-flow in an automated manner.

**Figure 1.**
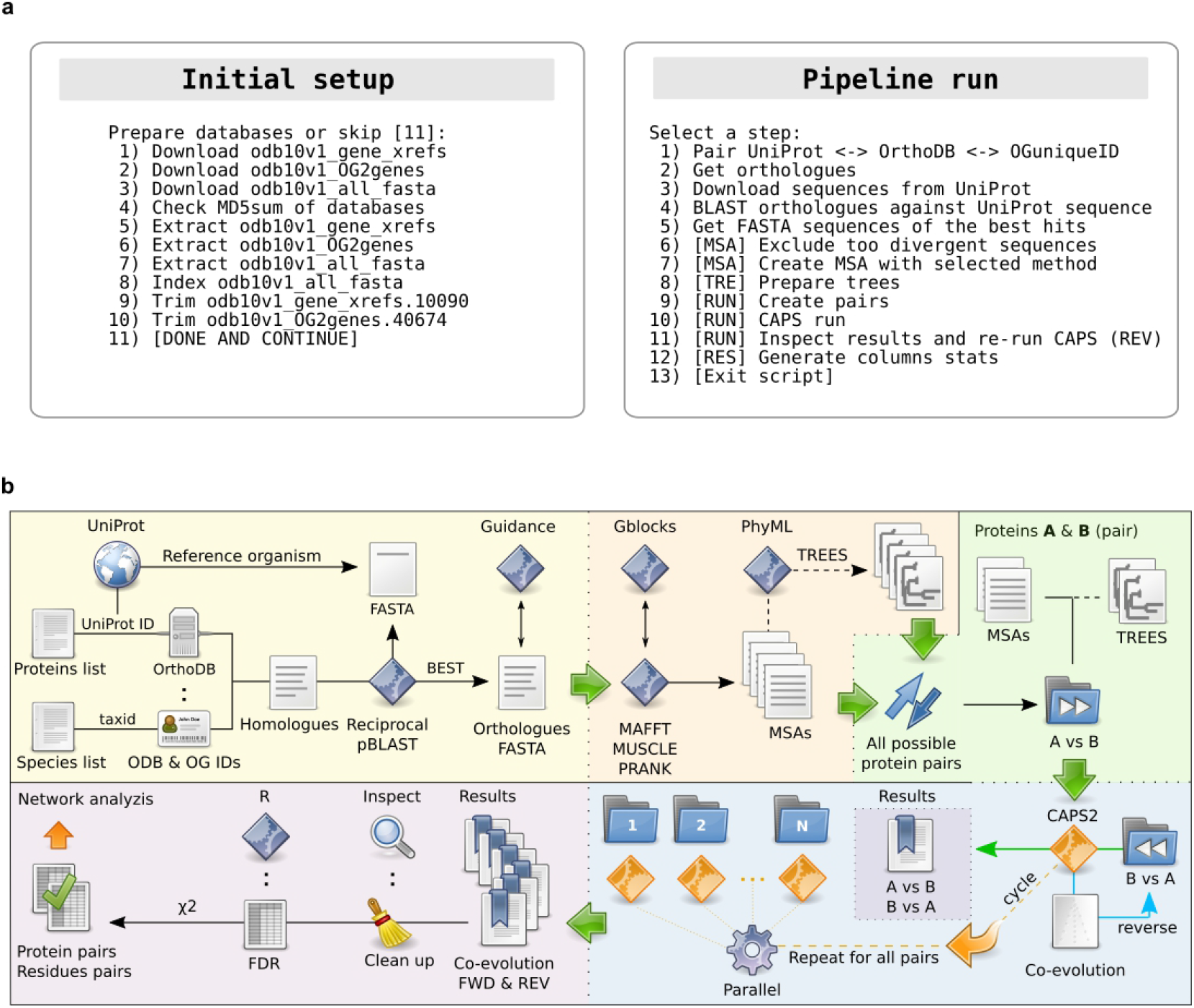
AutoCoEv pipeline. **A)** Menu overview. **Left**: Steps 1-7 download, check and extract databases; Steps 8-10 index and process the databases; **Right**: Steps 1-3 deal with homologous sequences retrieval; steps 4-5 carry out the identification of most appropriate orthologues; step 6 calls Guidance to exclude sequences that are too divergent; steps 7-9 create the MSA, phylogenetic trees and protein pairwise combinations; steps 10-11 run parallelized CAPS for each unique protein pair combination “bidirectionally”; step 12 processes results and does statistical analyses. **B)** Pipeline overview. **Yellow**. Reading the user-provided lists of proteins of interest and species to be searched, the script communicates between databases to extract genes (ODB) and orthologous groups (OG) identifiers (ID). Homologous sequences are then blasted against the UniProt sequences from the reference organism (e.g. mouse or human) in order to prepare a FASTA list of most appropriate orthologues. Before MSA, orthologues are assessed by Guidance and too divergent ones are removed; **Orange**. Orthologues are aligned by selected method (MAFFT, MUSCLE or PRANK) and scanned by Gblocks, to report regions of low quality. PhyML calculates trees from the MSA generated in the previous step, optionally using an external tree as a guide; **Green**. Create all unique protein pairs in folders, each folder having two sub-folders for MSA and (if prepared by PhyML) trees. **Blue**: CAPS2 is run for each protein pair folder in a parallelized fashion via GNU/Parallel If co-evolution is detected, CAPS2 is run again, this time “reversing” the protein load order (e.g. A vs B followed by B vs A). **Purple**. The output in each pairs folder is inspected and processed, followed by FDR correction of p-values and *Chi* squared test. Finally, the results are prepared as a table ready for the network analysis by Cytoscape.

As an input, AutoCoEv requires a list of proteins with their UniProt identifiers [19] and a list of species, for which orthologues will be searched. Optionally, a phylogenetic tree may be provided from an external source, such as TimeTree [20], to be used as a guide when trees are calculated from MSA (see later, *Multiple sequence alignments and trees*). Upon start, AutoCoEv offers to download the required databases from OrthoDB [21] and to run initial preparations, such as FASTA database indexing (Figure 1a, left). Once databases are in place and input files are loaded, the pipeline proceeds to the main menu that carries out the work-flow (Figure 1a, right).

### 2.2 Identification of orthologues

For each protein of in the user-provided list, AutoCoEv consults with OrthoDB, searching for homologues from the species of interest (Figure 1b, yellow panel). The script matches the UniProt ID of each protein to its OrthoDB ID, then extracts its unique orthologues group (OG) ID at a given level of organisms (e.g. *Eukaryota, Metazoa, Vertebrata, Tetrapoda, Mammalia*). This level, or node, is specified by the user and depends on the species for which orthologues are searched. The script will report proteins with missing OG IDs at OrthoDB, as well as, species for which no orthologue was found.

AutoCoEv prepares a list of homologues for each protein, however, there may be more than one per species, for example due to alternative splicing or gene duplication. Therefore, the homologues from each species are compared to the UniProt sequence of the user-provided protein, by pBLAST [22]. After this reciprocal BLAST against the “reference” organism, AutoCoEv selects the best hit per species. Importantly, users have the option to omit even the best hits if they do not pass certain criteria, such as identity to the reference sequence and alignment gaps. With this filtering step, the script avoids the inclusion of erroneous or not complete sequences that can skew the MSA in the next step. As a result, each protein holds a collection of automatically curated orthologous sequences, one per species. Before the MSA step (next), AutoCoEv additionally consults with program Guidance [23] to assess whether some orthologues are too divergent from the rest. The presence of such sequences can, again, affect the robustness of the alignment in the next step and it may be desirable to omit them.

### 2.3 Multiple sequence alignment and trees

CAPS2 detects co-evolution between two proteins by extrapolating from their MSAs, hence the quality of the alignments is of crucial importance [24]. For MSA creation (Figure 1b, orange panel), AutoCoEv offers a choice of three widely-used and accurate programs: MAFFT (Multiple Alignment using Fast Fourier Transform) [25], MUSCLE (MUltiple Sequence Comparison by Log-Expectation) [26] and PRANK (Probabilistic Alignment Kit) [27] (Figure 2B). Different MAFFT aliases are supported (e.g L-INS-i, E-INS-i, G-INS-i), while for PRANK an external phylogenetic tree (e.g. obtained from TimeTree) can be specified as a guide. After MSAs are generated, the script inspects them by program Gblocks [28], to assess the quality of the alignments regions. This information is reported in the final output, allowing the user to filter out co-evolving amino acids that belong to poorly aligned columns.

**Figure 2.**
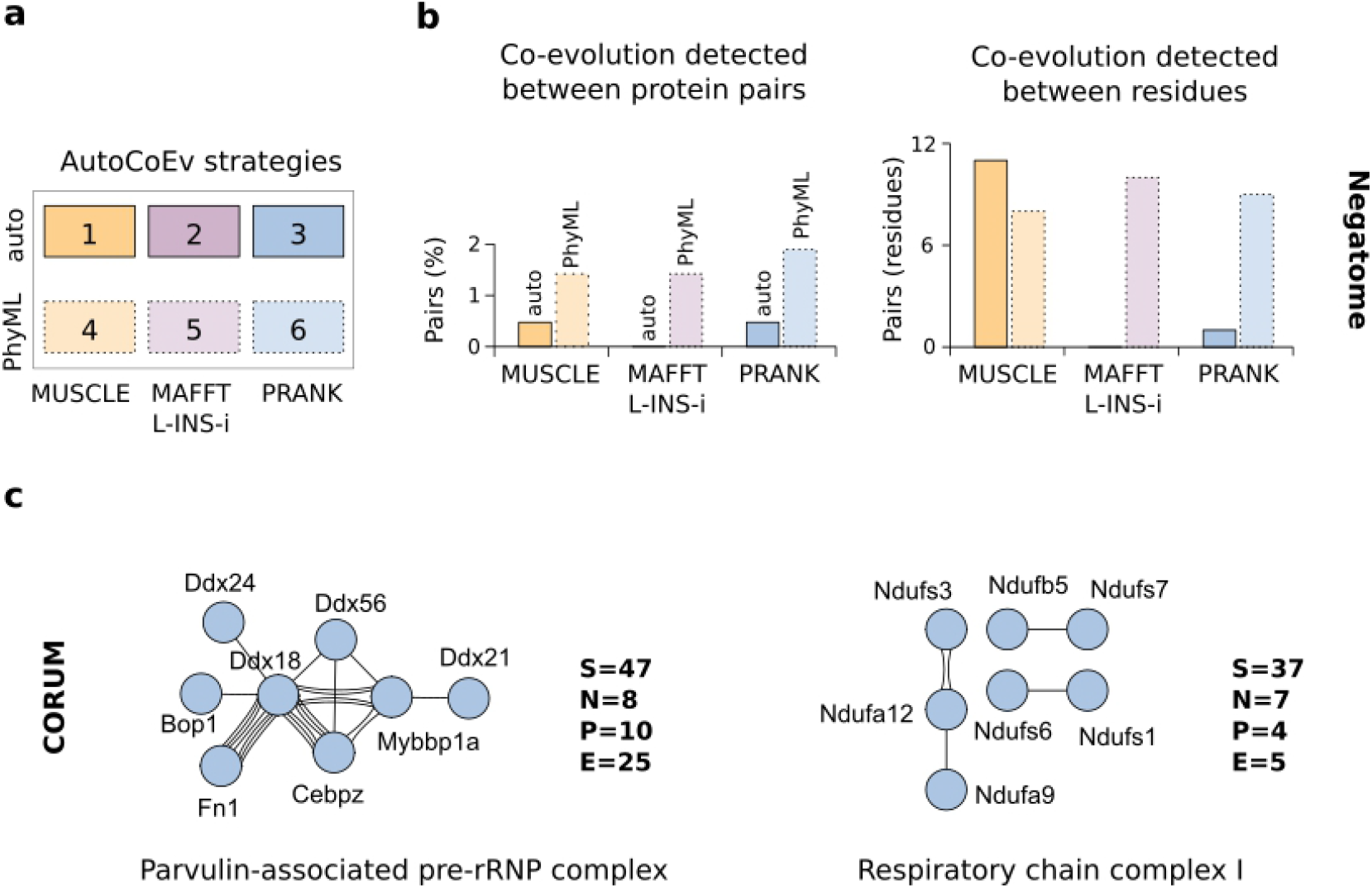
Determining AutoCoEv strategy. **A)** Strategies. Combinations of MSA method and phylogenetic trees calculation methods are referred as “strategies”. **B)** Negatome database. Percentage of protein pairs for which co-evolution was detected and the total number of co-evolving sites. C) CORUM database. Co-evolution was detected in 5 out of the first 10 complexes. Shown are the two largest. S: number of complex subunits (found in OrthoDB), P: protein pairs detected, E: number of edges (total number of co-evolving amino acids).

By default, CAPS2 generates its own BioNJ (neighbor-joining) distance-based phylogenetic trees from the proteins MSAs at runtime. The trees are not made available to the user, therefore we patched the program to print the tree to the output, allowing for inspection. Alternatively, trees calculated by another program can be used, which may improve the sensitivity of CAPS2. If this is preferred, AutoCoEv calls PhyML (Phylogenetic estimation using Maximum Likelihood) [29] (Figure 1b, orange panel). An external tree (e.g. from TimeTree) can be specified as a guide, while the generated trees can be rooted by TreeBeST [30], by minimizing height.

### 2.4 Detection of inter-protein co-evolution by CAPS2

The computational time required for the co-evolution detection presents a major bottleneck, as CAPS2 lacks CPU multi-threading. To overcome this limitation, AutoCoEv runs CAPS2 on individual protein pairs via GNU/Parallel [31], as described below.

First, AutoCoEv produces all unique pairwise combinations between the proteins from the user-provided list (Figure 1b, green panel). The script creates an individual folder dedicated to each pair and determines the species where an orthologous sequence was found for both proteins. Species that are not shared by the two proteins have their sequences removed from the MSAs by SeqKit [32], and are trimmed from the trees (if PhyML is used) by TreeBeST. This is important, since the presence of too many not shared species seems to deteriorate the stability of CAPS2. On the other hand, too little species result in poor reliability of the coevolution detection [11], therefore users can specify a minimum threshold of shared species for a protein pair (e.g. 20).

During the AutoCoEv development, we noticed that the order in which CAPS2 loads its input files, seems to have an effect on the inter-molecular analyses. Therefore, to improve the specificity and reliability of the analysis, we designed our script to run CAPS2 twice, so that those proteins pairs (e.g. A vs B) where co-evolution was detected in the first run get selected for asecond run, this time reversing the order (e.g. B vs A), by slightly renaming the files (Figure 1b, blue panel). Since CAPS2 loads input files randomly, we additionally patched the program to always load files in alphabetical order. Upon completion of the second run, AutoCoEv extracts the amino acid pairs predicted as co-evolving in both runs.

AutoCoEv does this for all protein pairs, using GNU/Parallel to spawn multiple instances of CAPS2, each operating in a single proteins pair folder. As a result, the script dramatically speeds up the time of computation.

### 2.5 Post-run processing of the results

At run time, CAPS2 uses an α-value threshold (e.g. *α* = 0.01) for the probability of error in rejecting the null hypothesis (type I error), when significant co-evolving sites are detected. Amino acid pairs that pass the threshold are reported in the results of CAPS2, however the actual p-values of their correlations are not. This poses limitations to statistical analyses, such as control of the false discovery rate (FDR), critical for large datasets. Therefore, we patched CAPS2 to calculate and output p-values when inter-protein co-evolution is searched (see Supplementary information, “*P-values of the results*”).

After CAPS2 runs are completed in all protein pair folders, AutoCoEv processes the results in several steps of filtering, sorting and assessment (Figure 1b, violet panel). Following initial clean-ups, the script calls R [33] to produce adjusted p-values of the co-evolving sites from each protein pair. By default, CAPS2 applies a chi squared (χ2) test when more than two proteins are analyzed, based on the number of detected co-evolving amino acids between them. In our pipeline, CAPS2 always runs for just 2 proteins at a time, therefore AutoCoEv replicates the χ^2^-test, when the analyses of all protein pairs are completed (see Supplementary information, *“Chi squared test”*).

Results are saved in two spreadsheet files: one containing all individual co-evolving amino acids, while the other summarizes the results per protein pair. Both spreadsheets are ready to import to Cytoscape [34] for network visualization and further analysis, such as filtering and cluster analysis.

## 3. Application

We searched for orthologues from 50 placental mammals (Table S1) in the analyses described below. The choice of methods for generation of the MSAs and phylogenetic trees is critical for evolutionary studies, therefore, we first tested 6 different combinations (Figure 2a). We used the Negatome database [35] of non-interacting proteins, for which we expect to detect little or no co-evolution.

We tested 211 protein pairs from mouse (Table S2) and observed that while all strategies predicted very low percentage of protein pairs with co-evolution, the strategies 2 and 3 (MSAs by MAFFT L-INS-i and PRANK with phylogenetic trees automatically generated by CAPS2), yielded the lowest numbers of co-evolving residues (Figure 2b, right). Using PhyML-generated phylogenetic trees in strategies 4-6 seemed to significantly increase the numbers of detected co-evolving amino acids. As a very low amount of coevolution was expected in this dataset and with the aim to minimize false positive hits, we decided to continue with the more conservative strategies with CAPS2-generated phylogenetic trees. In addition, the strategies 3 and 6 using PRANK showed the smallest fraction of protein pairs for which coevolution was detected in both CAPS2 runs of different directionalities but not on the same amino acids (therefore omitted in the final results), suggesting a better reliability with this MSA method (Supplementary figure S1a). To summarize, we considered that the best specificity was offered by strategy 3: combination of PRANK-generated MSAs with phylogenetic trees automatically calculated by CAPS2 at run-time.

We then used the CORUM database of known protein complexes [36], in order to test the selected strategy 3 on proteins for which co-evolution is expected. We analyzed the 10 largest complexes from mouse (number of subunits > 10), from which we detected co-evolution in 5 (Table S3, Figure 2c, Supplementary figure S1b). The highest number of co-evolving proteins were detected within the two largest complexes: the Parvulin-associated pre-rRNP complex and Respiratory chain complex. Although it is expected that such multiprotein complexes have gone through remarkable co-evolution, our analysis only assessed the situation in mammals and, thus, is likely to miss the interactions within highly conserved protein domains. Together the analysis of the negatome and the protein complexes pointed towards capabilities of AutoCoEv to predict protein-protein interactions and allowed us to proceed for large-scale data analysis.

### 3.1 Lipid rafts dataset

We used AutoCoEv to predict novel partners in a set of 324 proteins from mouse (Table S4), located at the lipid-raft membrane domains of B cells. The proteins were identified in a preceding study from our research group, by an APEX2 proximity biotinylation-based proteomics analysis [18].

Although our Negatome AutoCoEv runs inclined us to pick PRANK for our dataset, we again compared, in this large dataset, the MSAs produced by the other two programs, too. Mumsa [37], indicated that all three programs had produced high quality alignments, however the highest scores were clearly assigned to PRANK (Supplementary figure S2a). After screening for too divergent sequences and shared species, we had 46775 unique protein pairs to be tested for co-evolution. Following the CAPS2 double run step with PRANK alignments and automatic trees (strategy 3), we obtained a network of 61 nodes and 282 protein pairs (Figure 3a). The number of predicted co-evolving pairs was significantly lower than that when MUSCLE or MAFFT L-INS-i alignments were used. However, the results obtained with PRANK MSAs had best overlap with the results obtained by the other two methods (Supplementary figure S2b). The MSA quality scores and the best concordance of the co-evolving pairs with those detected by the other strategies (1 and 2), favored PRANK as the alignment method for subsequent analyses.

**Figure 3.**
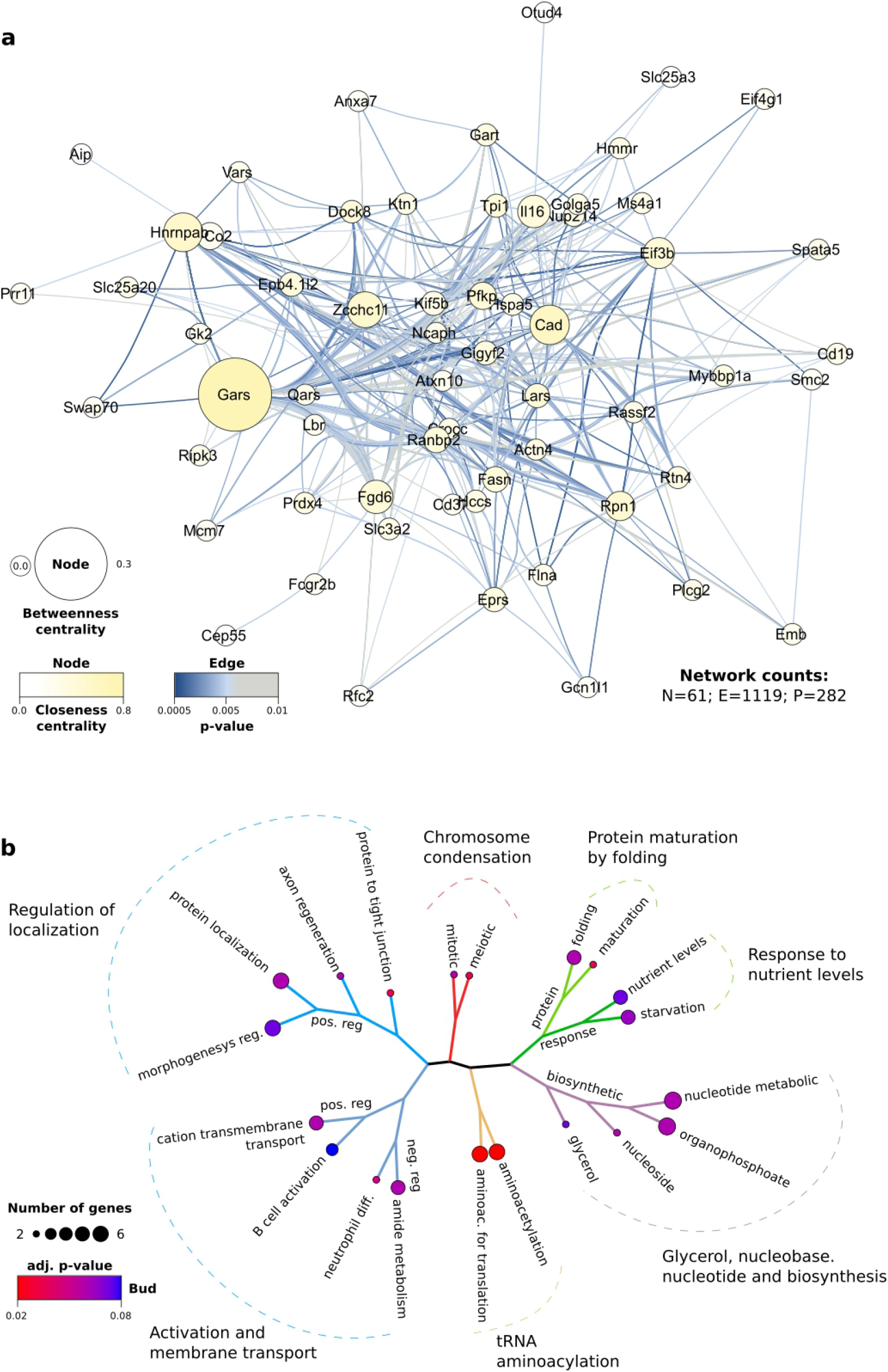
Proteins predicted to co-evolve in the dataset of 324 proteins from the vicinity of the B cell membrane rafts. **A)** Network analysis of the co-evolving pairs. Number of nodes (N), pairs (P) and edges (E) are indicated. Nodes size and colour reflect betweenness and closeness centrality, respectively. Edges colour corresponds to the co-evolution p-value of each amino acid pair. **B)** Gene ontology analysis of the network proteins. The cellular processes in which proteins are involved are indicated.

The network obtained from the predicted coevolving proteins had several major node hubs (Figure 3a), as defined by their closeness and betweenness centrality, namely Gars (Glycine-tRNA ligase), Cad (Carbamoyl-phosphate synthetase 2, Aspartate trans-carbamylase, and Dihydroorotase), Hnrnpab (Heterogeneous nuclear ribonucleoprotein A/B) and Eif3b (Eukaryotic translation initiation factor 3 subunit B). As reported by UniProt [38], all of them are multi-domain proteins, involved in translation, metabolism and transcription regulation. Performing gene ontology (GO) analyses on all 61 proteins indicated that they play a role in a wide range of processes and pathways (Figure 3b), such as in cell division, protein synthesis, cellular response, metabolism, membrane transport and more.

We sought to single out proteins that are tightly interlinked, in order to suggest potential candidates for further investigation in the wet-lab. We filtered the results by p-value (p < 0.005), alignment region quality (determined by Gblocks) and MSA column gaps (less than 20%), obtaining a smaller network, with an overall organization very similar to the parent (Figure 4a). About 25% of the protein pairs were also found in the STRING database (combined confidence score > 0.15), both before and after the network filtering (Figure 4b). Then, we preformed clustering analysis by CytoCluster (ClusterONE, number of nodes < 20) and distinguished a relatively compact cluster of 18 nodes (Figure 4c). The cluster incorporated total of 41 proteins pairs, 19 of which were found in STRING (Figure 4c).

**Figure 4.**
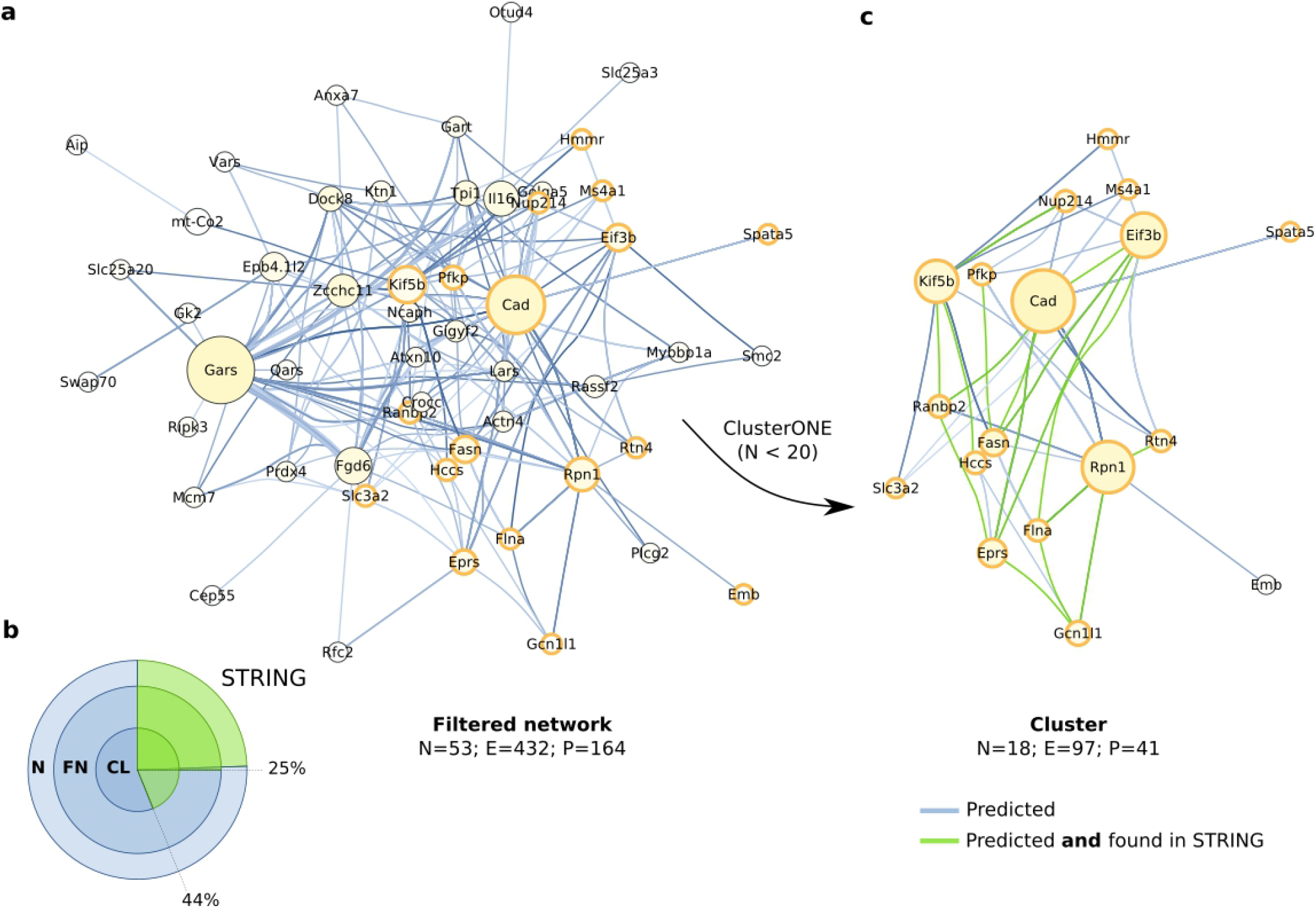
Cluster analysis of the co-evolution network. **A)** Cluster analysis of the co-evolution network. Edges were filtered by p-value (p<0.005), Gblocks score (good) and MSA column gaps (less than 20%). Nodes that are found in the cluster identified by ClusterONE (C, arrow) are outlined in orange. See Figure 3 legend for nodes size, nodes colour and edge colour. **B)** Protein pairs found in the STRING database. Overlapping circles represent the whole network (N), filtered network (FN, Figure 3a) and cluster (CL, Figure 4c). **C)** The identified cluster. Protein pairs supported also by STRING are shown in green.

The major hub node in the cluster is Cad, a large (243 kDa) protein with multi-catalytic activity, involved in de novo synthesis of pyrimidine [39]. Cad was predicted to co-evolve with 10 proteins (Supplementary figure S3a) via 7 amino acids: 20A, 1728L, 1887G, 1892A, 2108S, 2114S and 2160A. The residues were found in its GATase (glutamine amidotransferase), DHOase (dihydroorotase), DRBS (disordered region binding sites) and ATCase (Aspartate transcarbamylase) regions (Figure 5a).

**Figure 5.**
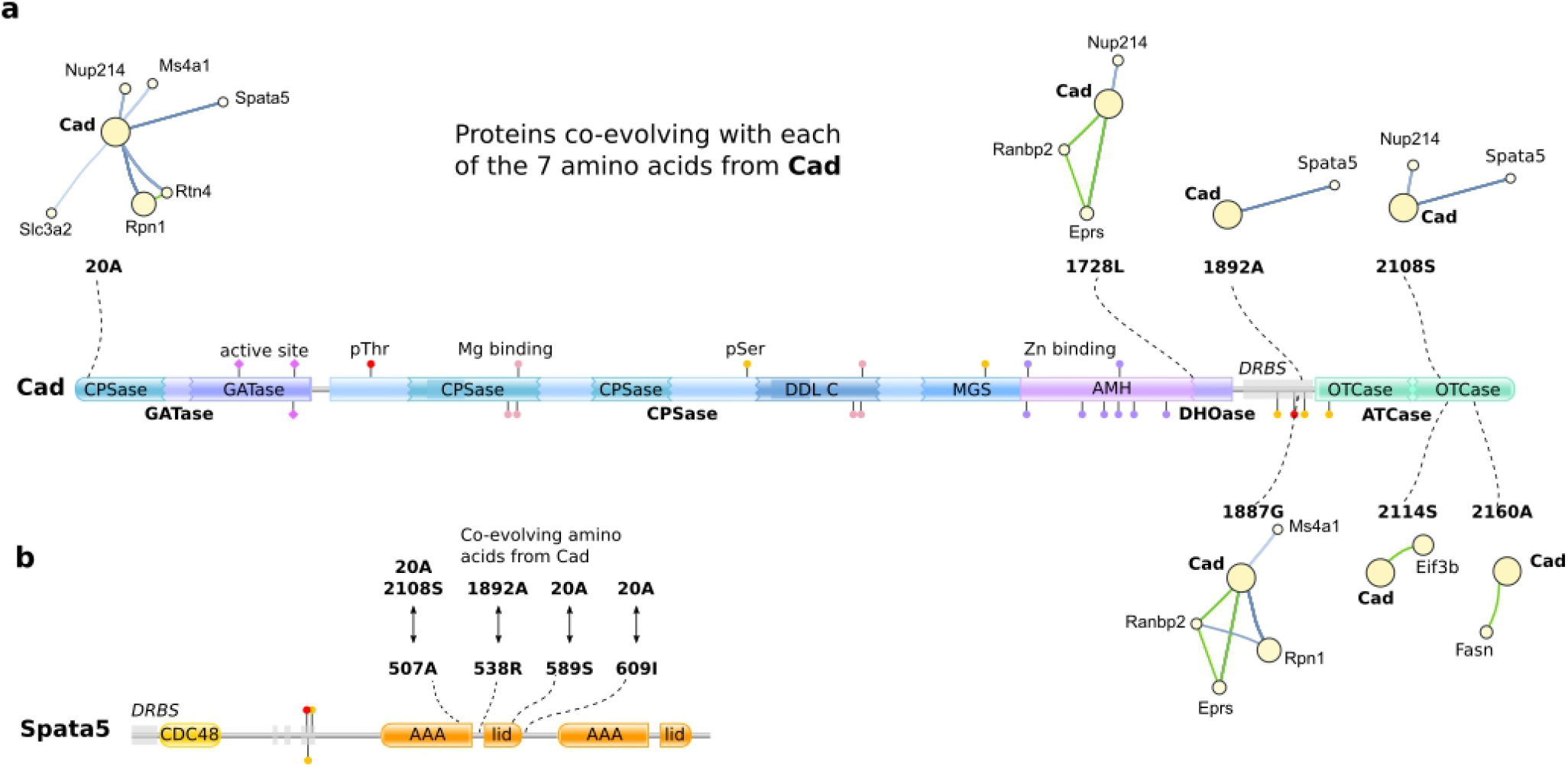
Suggested interactions between Cad and the proteins predicted to co-evolve with it. **A)** Co-evolving sites from Cad. The co-evolving amino acid residues are indicated by dashed lines and plotted onto the protein topology. Proteins co-evolving with each site are shown as mini-clusters. Protein regions are indicated in bold next to the scheme: GATase (glutamine amidotransferase), DHOase (dihydroorotase), DRBS (disordered region binding sites) and ATCase (Aspartate transcarbamylase); domains are indicated within scheme: CPSase (Carbamoyl-phosphate synthase small chain), DDL (D-alanine-D-alanine ligase), MGS (Methylglyoxal synthase), AMH (Amidohydrolase family domain), OTCase (Ornithine carbamoyltransferase, carbamoyl-P binding domain); active sites, phosphorylation sites (Ser, Thr) and metal-binding sites (Mn, Zn) are indicated as lollipops. **B)** The amino acid residues from Spata5 predicted to co-evolve with Cad. Amino acids are plotted onto the protein topology and paired sites from Cad are indicated. Domains: CDC48 (Cell division protein 48), AAA (ATPases associated with a variety of cellular activities), lid (AAA+ lid domain).

By GO analysis, Cad and its 10 co-evolving proteins were suggested to play roles in angiogenesis, ER, endothelium, membrane transport and translation (Supplementary figure S4). The proteins are predominantly cytoplasmic, nucleus-associated and membrane-associated (Table S5). The ATP-dependent chaperone Spata5 showed highest number of non-overlapping co-evolving residue pairs with Cad: 4 sites co-evolving with 3 sites from Cad (Supplementary figure S3b). They all resided within its first AAA–lid3 ATPase tandem domains, co-evolving with the CPSase domain, DRBS and the OTCase domain from Cad (Figure 5b).

## 4. Discussion

In this work, we present the AutoCoEv pipeline: an interactive script for the large-scale prediction of inter-protein co-evolution by CAPS2. Searching for signs of co-evolution in silico is a powerful means to predict novel functional interactions between proteins. By default, inter-molecular analysis in CAPS2 was designed for a handful of proteins, or a single pair, as illustrated by its web interface (http://caps.tcd.ie/caps/). However, the availability of CAPS2 for offline use, grants a great deal of flexibility achievable via scripting. Requiring only a list of proteins and a list of species, AutoCoEv automatically performs database searching and identification of orthologous sequences with a best hit. The pipeline offers further automation, parallelization and quality assessment on the subsequent steps and seamlessly achieves the batch processing of hundreds of input proteins. AutoCoEv also contains post-analysis tools enabling efficient analysis and ranking of the results. We propose that automated prediction of co-evolution provides a powerful and affordable tool to facilitate selection of candidates from large protein data sets for further analysis.

Prediction of co-evolution requires, at first, a collection of protein ortholoques from various species and generation of the MSAs. Already this is a sizable data mining and organizing task data, but is followed by extremely heavy residue-to-residue comparisons of MSAs. The challenging nature of the analysis is illustrated by the waiting times when using co-evolution analysis software that are available on servers. For instance, we used the BIS2, MISTIC, MISTIC2, ContactMap, GREMLIN and DCA servers aiming to analyze a single protein pair: Cad – Spata5. To detect inter-molecular co-evolution, BIS2 requires that the MSAs of the two proteins are concatenated and we did the same for MISTIC(2). The queue/runtime was ~22 hours for MISTIC, while MISTIC2, DCA and BIS2 crashed. GREMLIN and ContactMap did not accept MSA over 1000 and 1100 amino acids, while GREMLIN had the additional warning that 85 jobs were currently running and that our submission “may take forever to complete”. Thus, it is clear that large-scale analyses cannot be practically performed using the server-based tools accessible online. On contrary, AutoCoEv runs locally on Linux, thus avoiding queue waiting times and other limitations that arise when using a public server. The only results we obtained, from MISTIC, were very challenging to interpret, since the program (naturally) detected numerous inter-molecular co-evolving sites, making it virtually impossible to distinguish the inter-molecular co-evolution. For a comparison, running the CAPS2 bidirectional step via AutoCoev took ~48h on an i7-9700KF CPU (8 cores @3.6GHz) with 64GB RAM for 46775 protein pairs. In addition, CAPS2 does not have a MSA-length limitation (e.g. 1000 amino acids), allowing the processing of large proteins, such as Cad. The post-run processing by AutoCoEv provides users with a comprehensive table, that can also be directly imported in Cytoskape for further network analysis.

Since the choice of alignment software largely depends on the sequences being aligned, our script already drives three of the most widely used MSA programs [40], and we consider incorporation of additional methods in the future, such as T-coffee [41] and ClustalΩ [42]. For our dataset, PRANK appeared to be the most suitable MSA method, however users can choose also from MUSCLE or MAFFT. Using PhyML-calculated trees seemed to increase the sensitivity of CAPS2, something that was undesired in our analyses, as we aimed to minimize false-positives. However, if greater sensitivity is required or if co-evolution is initially expected, for example for known protein complexes, PhyML offers a reliable means for trees calculation outside of CAPS2. In addition, we are also planning to implement AutoCoEv wrappers around RAxML [43], MrBayes [44] and IQ-TREE [45] in the future.

While developing AutoCoEv, we applied several improvements to CAPS2 which, being an open source program, allows for feature implementation. Our patch making the program report p-values, greatly augments the verbosity of the results and allows for additional statistical tests. Moreover, allowing the user to inspect the phylogenetic trees produced by CAPS2 helps in the assessment of the results. Our observation that the order in which the two input files are loaded seems to matter for the outcome of the results was unexpected. Thus, to increase the confidence of the results, we opted to run the program twice, “bidirectionally”, then extract the residue pairs for which both runs agree. To ensure the repeatability of the process between different computers, we further patched CAPS2 to always sort input by alphabetical/numerical order. We believe this simple workaround greatly improved the specificity and the reliability of the program. Importantly, it should be noted that AutoCoEv is open-source and well amendable for inclusion of other programs, too. We welcome developers of co-evolution analysis programs to consider utilizing the AutoCoEv pipeline and testing their own program in the place of CAPS2, for high-throughput analysis.

Here, we analyzed hundreds of proteins on a regular computer, while the script is designed to work with even thousands of proteins, provided a high computing power is available. As an example, we focused on Cad, the central node within a compact cluster identified from our network. Cad, together with Eif3b, Fasn, Pfkp and Rnp, have been reported to localize in extracellular vesicles [46], supporting the predicted functional association between them. A novel relationship between Cad and Spata5 was suggested by their high number of co-evolving sites. Cad has been shown to locate towards mitochondria of mammalian spermatozoa [47], while Spata5, being an ATPase is essential in mitochondrial morphogenesis during early spermatogenesis [48]. Therefore, we hypothesis that Cad may provide a pyrimidine nucleotide pool that could stimulate Spata5’s ATPase during spermatogenesis. The proteins discussed here represent only a nominal part of the full myriad of possible candidates for further detailed analyses.

We trust that AutoCoEv, as an affordable and unbiased *in silico* analysis, could benefit various large scale protein interaction studies, like imaging mass spectrometry [49], by providing another viewpoint to the connections between the proteins and, thus, help in identifying interesting proteins or pathways for further studies.

## 4. Materials and methods

### 4.1 Availability and required software

AutoCoEv is written in BASH and is under MIT license, freely available from the GitHub repository of our group (https://github.com/mattilalab/autocoev). Development was done on Slackware (http://www.slackware.com/) and CRUX (https://crux.nu/) distributions of GNU/Linux.

Software tools that AutoCoEv drives and their versions used in our analyses are: CAPS (2.0 patched, see Supplementary Information), Datamash (1.7), Exonerate (2.4.0), Gblocks (0.91b), Guidance (2.02), MAFFT (7.471), MUSCLE (3.8.1551), NCBI BLAST+ (2.12.0), PRANK (170427), Parallel (20211122), PhyML (3.3.20200621), R (4.1.2), SeqKit (0.16.1), squizz (0.99d) and TreeBeST (git:347fa82, Ensembl modifications).

See AutoCoEv on GitHub for details, manual, as well as for instructions for setting up the pipeline on Ubuntu (https://ubuntu.com/) or Debian (https://www.debian.org/). We provide a pre-compiled, static binary of CAPS2 with our patches applied, and a virtual machine image with all requirements pre-installed.

### 4.2 Databases

AutoCoEv uses the following databases from OrthoDB (https://www.orthodb.org/): odb10v1_all_fasta, odb10v1_gene_xrefs, odb10v1_OG2genes. The script also communicates with UniProt (https://www.uniprot.org/) to download the latest sequence of each protein of interest.

Non-interacting protein pairs from mouse (Table S2) were obtained from Negatome (http://mips.helmholtz-muenchen.de/proj/ppi/negatome/). Conversely, protein complexes were obtained from CORUM (http://mips.helmholtz-muenchen.de/corum/) and sorted by size. The first 10 largest (subunits > 10) from mouse were retrieved (Ids: 3047, 382, 39, 6938, 538, 2750, 496, 572, 582, 1001), whereas co-evolution was found within 5 complexes (Table S3).

Protein networks were compared against STRING (https://string-db.org/).

### 4.3 Proximity biotinylation

For details, see Awoniyi et al bioRxiv [18], a preceding study from our group, published in parallel with this work. Briefly, lysates of B cells stimulated with 10μg/mL antibody against BCR were collected after 0 min, 5 min, 10 min and 15 min time points. The APEX2 system was used to induce biotinylation of proteins within 20 nm range in close proximity to the BCR. Samples were subjected to streptavidin affinity purification followed by mass spectrometry analysis. MaxQuant (1.6.17.0) was used for database search and after differential enrichment analysis with NormalyzerDE (1.6.0), a list of 346 proteins, proposed as raft-resident, was prepared. From these proteins, 324 were found in OrthoDB and used with here with AutoCoEv.

### 4.4 AutoCoEv run-time parameters

In our analyses, we used 50 mammalian species (Table S1) and configured Mus musculus, as a reference organism (taxid: 10090) and Mammalia as OrthoDB node level (taxid: 40674) in settings.conf. For the reciprocal BLAST, we set the minimum allowed alignment identity to 35% and the maximum allowed gaps to 25%. Guidance was used with MUSCLE and had a cutoff of 0.95. Three MSA methods were used in independent runs: MUSCLE, MAFFT alias L-INS-I and PRANK, while Gblocks allowed gaps were set to half (-b5=h). When PhyML trees were used, PhyML was run with default settings and the produced trees were rooted by TreeBeST. At the protein pairing step, the minimum required common species between each two proteins was set to 20. CAPS2 was run with bootstrap threshold 0.6 and convergence option (-c).

### 4.5 Protein Characterization

Protein sequence functional information was retrieved using UniProt. Domain organizations was searched at SMART [50] (Simple Modular Architecture Research Tool, http://smart.embl-heidelberg.de/) and Pfam [51] (http://pfam.xfam.org/). Disordered region binding sites were predicted by Anchor2/IUPred3 [52] (https://iupred3.elte.hu/). Graphical representation of proteins was rendered using Pfam cusrtom domain generator (http://pfam.xfam.org/generate_graphic/).

### 4.6 Data analyses

Post-run analyses were done by R (https://www.r-project.org/) and Gnumeric spreadsheet (http://www.gnumeric.org/). Gene Ontology was done by clusterProfiler (4.2.2) package for R [53]. Venn diagrams were generated by DeepVenn [54]. Networks were visualized in Cytoscape (3.8.2) and cluster analyses were performed by CytoCluster/ClusterONE [55]. All figures were assembled in Inkscape (https://inkscape.org/) while icons artwork in Figure 1 is from the Tango icon project (http://tango.freedesktop.org/).

## Supplementary materials

The following supporting information is included in the end of this document: Supplementary figures S1-S4; Supplementary tables S1-S5; Supplementary information.

## Author contributions

PBP developed AutoCoEv, analyzed the data, designed the figures and wrote the manuscript. LOA developed the statistical evaluation of the results by R scripting and performed GO analyses. VŠ conceived the original idea of the automated pipeline. MÖB performed proteins characterization and analyses. PKM contributed to the study design, trouble-shooting, and manuscript preparation.

## Institutional Review Board Statement

Not applicable.

## Informed Consent Statement

Not applicable.

## Data Availability Statement

AutoCoEv is available at https://github.com/mattilalab/autocoev

## Acknowledgements

We thank the members of the Lymphocyte Cytoskeleton lab for their critical comments on our manuscript. The authors would like to thank Martti Tolvanen for discussions on the theoretical basis and editing the final manuscript, Akseli Mantila for help with the C++ code of CAPS2 and Dian Dimitrov for advising on mathematics and statistics.

