## Supplementary for "AutoCoEv – a high-throughput *in silico* pipeline for predicting inter-protein co-evolution"

#### Supplementary figures

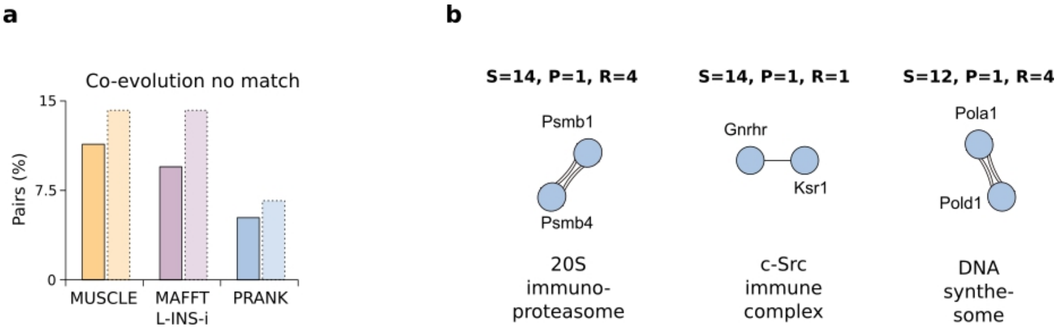

**Supplementary figure S1.** Determining AutoCoEv strategy. **A)** Non-matched co-evolution. Protein pairs for which co-evolution was detected in both CAPS2 runs, but not on the same amino acids. **B)** CORUM database. Co-evolution was detected in 5 out of the 10 largest complexes. Shown are the second 3.

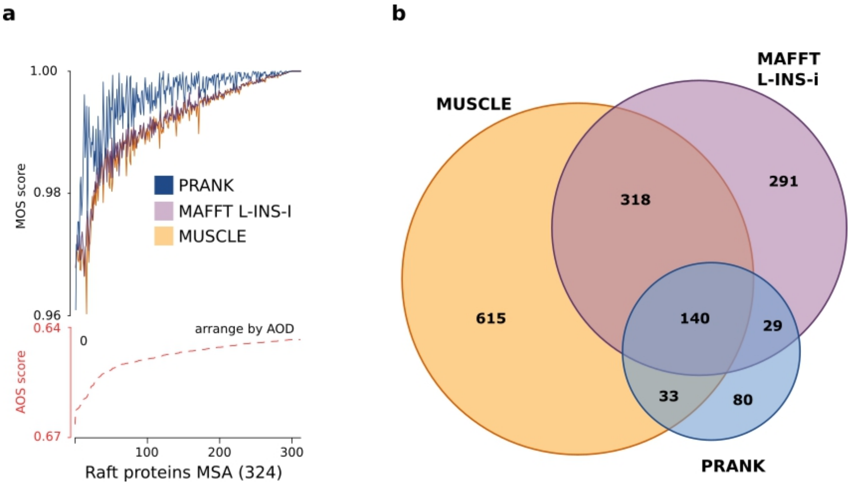

**Supplementary figure S2.** Assessing the alignment methods for the large-scale protein dataset. **A)** The quality of the MSAs. Alignments created by MUSCLE, MAFFT L-INS-i and PRANK were compared by Mumsa and sorted by their average overlap score (AOS), which reflects the difficulty of aligning the sequences. The multiple overlap score (MOS), which indicates the quality of each individual alignment was plotted for the three methods. **B)** Comparison of protein pairs detected to co-evolve. Overlap of the protein pairs for which co-evolution was detected from alignments created by MUSCLE, MAFFT L-INS-i and PRANK is indicated as a Venn diagram.

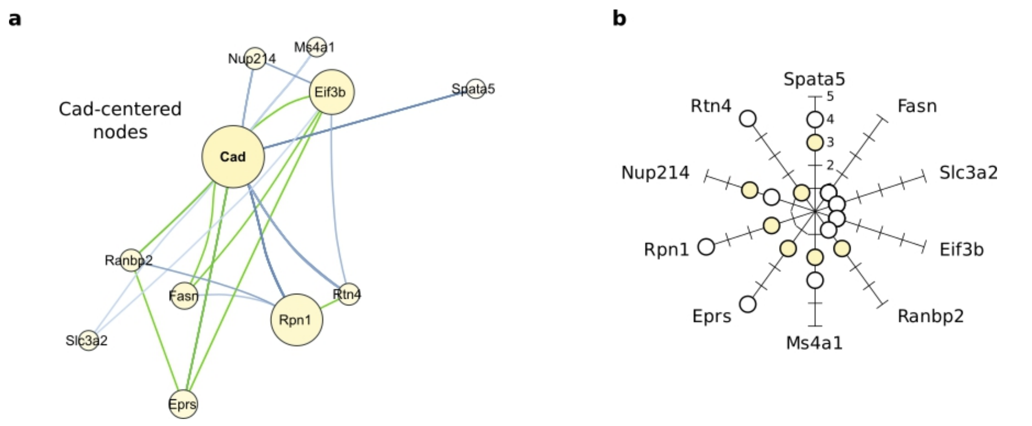

**Supplementary Figure S3.** Proteins co-evolving with Cad. **A)** Cad-centered proteins. Protein pairs supported also by STRING are shown in green. **B)** Number of co-evolving sites between Cad and its interacting proteins. The numbers of co-evolving sites between Cad (yellow circle) and the proteins predicted to co-evolve with it (whitecircle) are indicated.

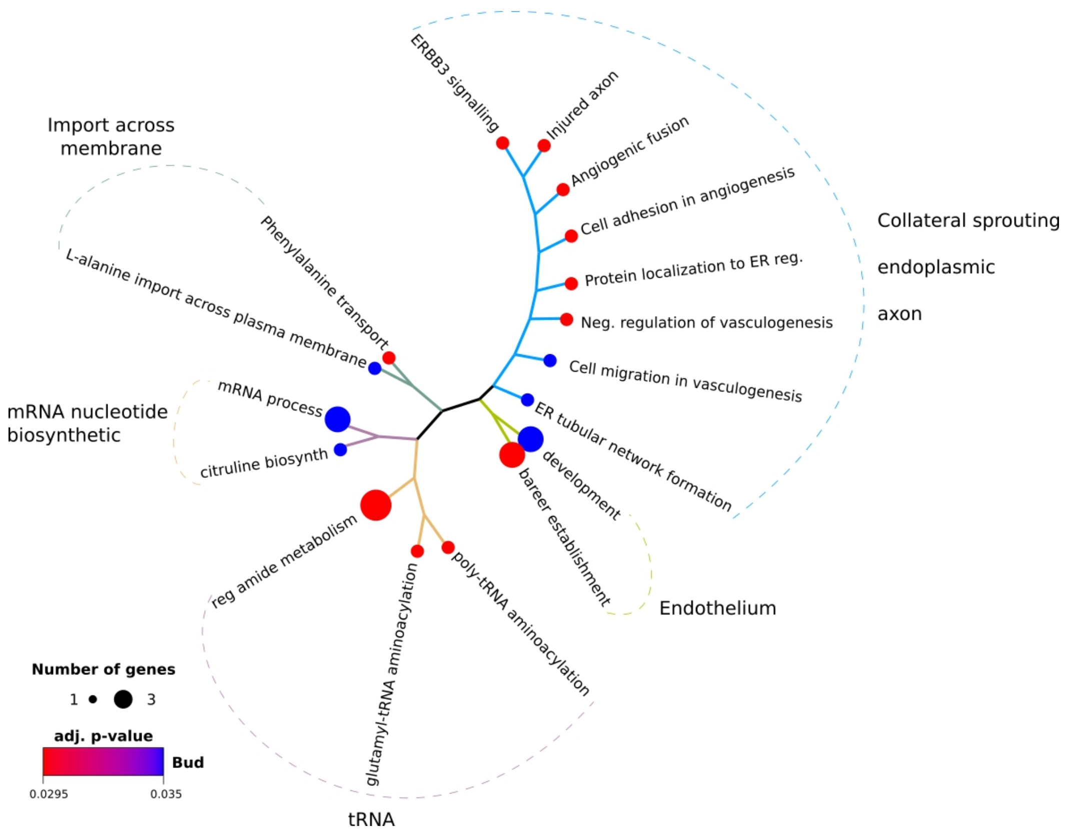

**Supplementary figure S4.** Gene ontology analysis of the clustered proteins. The cellular processes in which the proteins are involved are indicated.

### Supplementary information

**P-values of the results**

At run time, CAPS2 sets an $\alpha$-value threshold (e.g. $\alpha$ = 0.01) for the probability of error when rejecting the null hypothesis (type I error), when significant co-evolving sites are detected. Results with a probability of type I error below $\alpha$ are reported, however their actual p-values are not, which poses limitations to rank or compare the data between protein pairs. Therefore, we patched CAPS2 to output p-values when inter-protein co-evolution is searched, following the run steps of the program.

First, based on the input data, CAPS2 simulates a number of random MSA pairs, and tests all sites combinations for co-evolutionary correlation. The acquired correlation values are stored in a vector, the size of which ($V_{SIZE}$) depends on the MSA lengths of the two proteins ($L_1$ and $L_2$), and the number of simulations (r) performed (Equation 1). The correlations are sorted by value (from 0 to 1) and the vector hereafter serves as a null data distribution to which the real data will be compared. The $\alpha$-value derives an index number ($I_{THRESH}$) within the vector (Equation 2), whereas the value contained at the indexed position is set as a correlation threshold ($C_{THRESH}$) (Figure S1).

(1) $V_{SIZE} = L_1 \times L_2 \times r$

(2) $I_{THRESH} = V_{SIZE}(1 - \alpha) + 1$

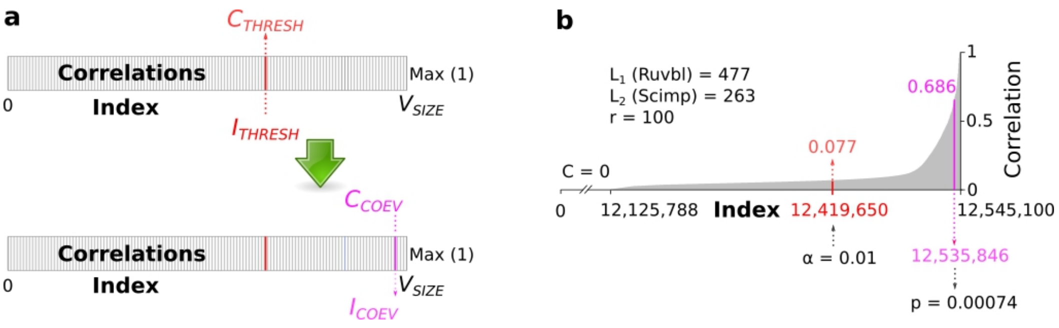

**SI Figure 1. Determining p-values.** A) Threshold and correlation. Correlation values of simulated data are stored in a vector (rectangular box) and sorted by size from 0 to 1 (Max). The vector size ($V_{SIZE}$) reflects the total number of possible correlations multiplied by the number of simulations. The threshold position within the vector (index, $I_{THRESH}$) is determined by α in Equation 1, and its correlation value ($C_{THRESH}$) is set as a cut-off for the correlations detected in the user-provided data ($C_{COEV}$). With our patch, CAPS2 ranks the position of each detected correlation value within the vector, determines its index ($I_{COEV}$) and uses it in Equation 2 to calculate the corresponding p-value. B) Example data. Protein pair Ruvb1 and Scimp was analysed in our preliminary runs. The lengths of Ruvb1 and Scimp MSAs (L1 and L2) and the number of simulations (r) are reflected in $V_{SIZE}$ = 12,545,100. The contents of the vector holding correlation values (C) from the simulated data are shown: C = 0 (no correlation) in blue (X-axis) and C > 0 in grey, starting at index 12,125,788. The threshold value position (red) is determined by α in Equation 1 and returns $C_{THRESH}$ = 0.077. A pair of co-evolving with a $C_{COEV}$ = 0.686, was ranked within the vector at index 12,535,846; the index was in turn used in Equation 3 to calculate p = 0.00074 for the correlation.

The threshold is thus specific for each protein pair, and correlations detected from the real data, that rank above it are deemed as being significant. With our patch (see "Patch for verbose output" for detailed information), after coevolution has been detected between two sites, CAPS2 ranks its value within the vector and determines the corresponding index ($I_{COEV}$). After "searching back" the $I_{COEV}$ (SI Figure 1a), CAPS2 can calculate the corresponding p-value (Equation 3). An actual example is shown in SI Figure 1b.

$$p = \frac{(1 + V_{SIZE} - I_{COEV})}{V_{SIZE}} \quad (3)$$

When analysing a protein pair, CAPS2 estimates the correlation between two sites bidirectionally (e.g. for sites A and B, correlation is estimated: A → B and A ← B), exporting their mean value in the results (this is not to be confused with the bidirectional run of CAPS2 that AutoCoEv performs, by loading the two MSAs in alternating order.). On rare occasions, CAPS2 assigns a negative value at the bidirectional correlation estimation step. Since this usually yields a mean value lower than the threshold, AutoCoEv dismisses site pairs where a negative correlation was estimated.

**Patch for verbose CAPS2 output**

Our caps_verbose.patch introduces two modifications to CAPS2, making the program produce more verbose output. First, the program will use *TreeTemplateTools* from bpp-phyl to export CAPS2 generated trees by *treeToParenthesis*, if none were supplied by the user at run-time.

When phylogenetic trees are not supplied by the user, CAPS generates its own trees automatically:

```
tree1 = create_input_tree(vec1.names, vec1.sequences);
tree2 = create_input_tree(vec2.names, vec2.sequences);
```

Our patch simply outputs the trees by *treeToParenthesis*:

```
string temptre1 = TreeTemplateTools::treeToParenthesis(*tree1, true);
string temptre2 = TreeTemplateTools::treeToParenthesis(*tree2, true);
```

*P-values output*

CAPS simulates number (-r) of random MSA, where each simulated alignment has the same number of columns (length) as the real data and is tested by the same method. This ensures that the data is compared to a *null* distribution without coevolution pressures. Correlation information from the simulated data is stored and sorted into vector *totaltemp*. The correlation *threshold* for each protein pair is based on the simulated data in *totaltemp*, found at a certain index, *value*. It is determined by the alpha (-a) run-time option, referred here as *threshval*:

```
[1] int value = floor(((totaltemp.size())*(1-(threshval))))+1;
threshold = totaltemp[value];
```

Correlation between residues R1 and R2 is determined as the mean of correlation R1→R2 and correlation R2→R1. Each must be higher than the *threshold*, whereas *thresholdR* is always 0.01:

```
if((fabs(Correl1[cor])>=threshold && fabs(Correl1[cor])>=thresholdR)
&& (fabs(Correl2[cor])>=threshold && fabs(Correl2[cor])>=thresholdR))
```

After coevolution between sites has been determined, CAPS2 ranks the correlation value within totaltemp vector containing the null simulated data. We seek to calculate the p-value of each correlation using the formula from [1], where *value* is the correlation closest *index* within the *totaltemp* vector. We introduce a new function *getIndex*, which ranks a value *K* within a vector *v*, defines its lower bound value as *it*, then determines the *index* of *it* within the vector. Here, *v* = *totaltemp* and *K* = Correl_[cor] and the p-value of Correl_[cor] is returned as *alphathresh*:

```
double getIndex(std::vector<double> const& v, double K) {
    auto const it = std::lower_bound(v.begin(), v.end(), fabs(K));
      if (it != v.end()) {
        int index = distance(v.begin(), it);
alphathresh = (((int)1+(double)v.size()-(int)index)/(double)v.size());
return alphathresh;
      }
      else {
        cerr << "ELEMENT NOT FOUND!" << endl;
      }
}
```

We use the index of lower bound *it*, since an exact match of the correlation value Correl_[cor] is unlikely to be found within *totaltemp*:

```
double Pvalue1 = getIndex(totaltemp, Correl1[cor]);
double Pvalue2 = getIndex(totaltemp, Correl2[cor]);
```

##### Chi squared test

We attempted to replicate in AutoCoEv the $\chi^2$ test performed by CAPS2 when more than 2 proteins are evaluated. For each single protein (hereby referred as Protein *0*) the number of co-evolving amino acid pairs detected with each other protein are considered (SI Figure 2), as follows:

**x:** number of co-evolving amino acid pairs between Protein *0* and Protein *i*
**t**: total number of possible amino acid pairs combinations (eq. 4) between Protein 0 and Protein *i*

(4) $t_i = L_0 \times L_1$ , see eq. 1

**b**: background co-evolution for Protein 0 against all other proteins

(5) $$b = \frac{\sum_{i=1}^{n} \frac{x_i}{t_i}}{n}$$

**e**: expected number of co-evolving pairs for Protein 0 and Protein i

$$e_i = bt_i$$

Co-evolution between Protein 0 and Protein *i* passes the test if:

(7) $$\chi^2 = \frac{(x_i - e_i)^2}{e_i}$$

$x_i > e_i$

$x_i > 3.84146$ ($\chi^2 = 0.05$)

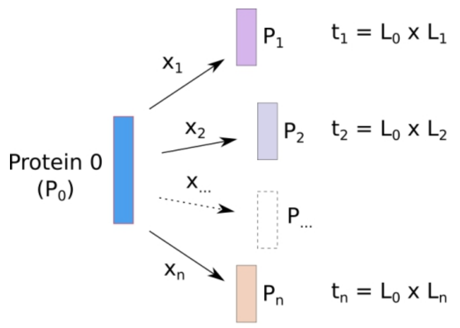

$$b = \frac{\left(\frac{x_1}{t_1} + \frac{x_2}{t_2} + \cdots + \frac{x_n}{t_n}\right)}{n}$$

**SI Figure 2.** Co-evolution between Protein 0 and all other proteins (n). Number of co-evolving amino acid pairs (x), total number of possible amino acid combinations (t) between $P_0$ and each of the other proteins ($P_1$, $P_2$, ..., $P_n$) are indicated. Equation shows calculation of background (b), following eq. 5.

##### Correlation normalized to threshold

Since correlation values ($C_{COEV}$) derived by CAPS2 cannot be directly compared between protein pairs, AutoCoEv calculates normalized values to the threshold ($C_{THRESH}$) of each protein pair, where 1 is the maximum possible correlation value:

(8) $$C_{NORM} = \frac{(C-T)}{(1-T)}$$

### Supplementary tables

**Table S1. Mammalian species used for AutoCoEv analysis.**

| Species 1-17 | | Species 18-34 | | Species 35-50 | |
|---|---|---|---|---|---|
| **taxid** | **Latin name** | **taxid** | **Latin name** | **taxid** | **Latin name** |
| 9606 | Homo_sapiens | 10116 | Rattus_norvegicus | 9796 | Equus_caballus |
| 9598 | Pan_troglodytes | 10029 | Cricetulus_griseus | 9807 | Ceratotherium_simum |
| 9597 | Pan_paniscus | 10181 | Heterocephalus_glaber | 9615 | Canis_lupus |
| 9593 | Gorilla_gorilla | 10141 | Cavia_porcellus | 9669 | Mustela_putorius |
| 9601 | Pongo_abelii | 43179 | Ictidomys_tridecemlineatus | 9646 | Ailuropoda_melanoleuca |
| 61853 | Nomascus_leucogenys | 9986 | Oryctolagus_cuniculus | 9685 | Felis_catus |
| 60711 | Chlorocebus_sabaeus | 9978 | Ochotona_princeps | 132908 | Pteropus_vampyrus |
| 9541 | Macaca_fascicularis | 482537 | Galeopterus_variegatus | 59463 | Myotis_lucifugus |
| 9544 | Macaca_mulatta | 9395 | Tupaia_glis | 143292 | Manis_pentadactyla |
| 9555 | Papio_anubis | 9365 | Erinaceus_europaeus | 9783 | Elephas_maximus |
| 43780 | Nasalis_larvatus | 42254 | Sorex_araneus | 9785 | Loxodonta_africana |
| 61622 | Rhinopithecus_roxellana | 9823 | Sus_scrofa | 99490 | Loxodonta_cyclotis |
| 27679 | Saimiri_boliviensis | 9913 | Bos_taurus | 9813 | Procavia_capensis |
| 1868482 | Carlito_syrichta | 9940 | Ovis_aries | 9371 | Echinops_telfairi |
| 30608 | Microcebus_murinus | 9739 | Tursiops_truncatus | 9361 | Dasypus_novemcinctus |
| 30611 | Otolemur_garnettii | 9767 | Balaenoptera_acutorostrata | 9358 | Choloepus_hoffmanni |
| 10090 | Mus_musculus | 30538 | Vicugna_pacos | | |

**Table S2. Negatome database protein pairs from mouse (*Mus musculus*)**

| Prot. A | Prot. B | PubMed ID | Evidence |
|---|---|---|---|
| O08599 | P70452 | 16186111 | MI:0007 - anti tag coimmunoprecipitation |
| O08599 | Q80T23 | 16186111 | MI:0007 - anti tag coimmunoprecipitation |
| O08599 | Q8K1E0 | 16186111 | MI:0007 - anti tag coimmunoprecipitation |
| O08599 | Q99N48 | 16186111 | MI:0007 - anti tag coimmunoprecipitation |
| O08599 | Q99N50 | 16186111 | MI:0007 - anti tag coimmunoprecipitation |
| O08674 | O35516 | 9111338 | MI:0059 - gst pull down |
| O08674 | P31695 | 9111338 | MI:0059 - gst pull down |
| O08674 | Q01705 | 9111338 | MI:0059 - gst pull down |
| O08674 | Q61982 | 9111338 | MI:0059 - gst pull down |
| O08715 | P31324 | 11742813 | MI:1200 - immunocytochemistry |
| O08849 | Q9Z0G0 | 9770488 | MI:0018 - two hybrid |
| O08915 | P53762 | 9083006 | MI:0019 - coimmunoprecipitation |
| O08983 | Q99KG7 | 12663659 | MI:0018 - two hybrid |
| O35551 | Q3UMR0 | 16525121 | MI:0019 - coimmunoprecipitation |
| O35625 | P70339 | 10428961 | MI:0007 - anti tag coimmunoprecipitation |
| O35657 | P01132 | 2690595 | MI:0004 - affinity chromatography technology, MI:0110 - text mining |
| O35717 | P05532 | 14707129 | MI:0019 - coimmunoprecipitation |
| O35718 | P05532 | 14707129 | MI:0019 - coimmunoprecipitation |
| O35718 | Q62120 | 10534114 | MI:0019 - coimmunoprecipitation, MI:0110 - text mining |
| O35984 | Q7TPV4 | 17875935 | MI:0019 - coimmunoprecipitation |
| O55003 | Q99JI6 | 17928295 | MI:0018 - two hybrid |
| O55131 | P42208 | 16914559 | MI:0055 - fluorescent resonance energy transfer |
| O70355 | P09793 | 17237401 | MI:0054 - fluorescence-activated cell sorting |
| O70355 | P31041 | 17237401 | MI:0054 - fluorescence-activated cell sorting |
| O70355 | Q7TSA3 | 17237401 | MI:0054 - fluorescence-activated cell sorting |
| O70355 | Q9WVS0 | 17237401 | MI:0054 - fluorescence-activated cell sorting |
| O70433 | Q62219 | 16737959 | MI:0019 - coimmunoprecipitation |
| O70622 | Q7M6Z0 | 17189258 | MI:0415 - enzymatic studies |
| O70622 | Q8K0S5 | 17189258 | MI:0415 - enzymatic studies |
| O88346 | P0C605 | 10671526 | MI:0018 - two hybrid |
| P01582 | Q61730 | 9620655 | MI:0030 - cross-linking study, MI:0110 - text mining |
| P02340 | Q6I6G8 | 12890487 | MI:0019 - coimmunoprecipitation |
| P03995 | P05627 | 15576676 | MI:0096 - pull down; MI:0019 |
| P05627 | P28738 | 15576676 | MI:0096 - pull down; MI:0019 |
| P05627 | P33175 | 15576676 | MI:0096 - pull down; MI:0019 |
| P05627 | P57776 | 15576676 | MI:0096 - pull down; MI:0019 |
| P05627 | Q61768 | 15576676 | MI:0096 - pull down; MI:0019 |
| P05627 | Q6PER3 | 15576676 | MI:0096 - pull down; MI:0019 |
| P05627 | Q9ESN9 | 15576676 | MI:0096 - pull down; MI:0019 |
| P06876 | P02340 | 8616838 | MI:0019 - coimmunoprecipitation, MI:0070 - mobility shift |
| P08752 | Q76MZ3 | 15525651 | MI:0059 - gst pull down |
| P09803 | Q8BN78 | 10207085 | MI:0019 - coimmunoprecipitation |
| P13405 | Q61501 | 7675450 | MI:0413 - electrophoretic mobility shift assay |

| | | | |
|---|---|---|---|
| P13808 | Q02357 | 8798415 | MI:0019 - coimmunoprecipitation |
| P15499 | Q9JJZ9 | 20238003 | MI:0019 - coimmunoprecipitation |
| P16054 | Q91YM2 | 14636894 | MI:0019 - coimmunoprecipitation |
| P17679 | Q80YQ2 | 17132730 | MI:0096 - pull down |
| P17679 | Q924H2 | 17132730 | MI:0096 - pull down |
| P17679 | Q9CQA5 | 17132730 | MI:0096 - pull down |
| P17679 | Q9CZB6 | 17132730 | MI:0096 - pull down |
| P17679 | Q9D7W5 | 17132730 | MI:0096 - pull down |
| P17918 | Q9ESU6 | 12192049 | MI:0006 - anti bait coimmunoprecipitation |
| P20917 | P11276 | 2446864 | MI:0892 - solid phase assay |
| P20917 | P13595 | 2446864 | MI:0892 - solid phase assay |
| P20917 | P20917 | 2446864 | MI:0892 - solid phase assay |
| P20917 | Q8K0S5 | 17189258 | MI:0415 - enzymatic studies |
| P21183 | P04401 | 1942734 | MI:0004 - affinity chromatography technologies |
| P23804 | P02340 | 10075719 | MI:0019 - coimmunoprecipitation |
| P26450 | P97465 | 18204460 | MI:0019 - coimmunoprecipitation |
| P26955 | P04351 | 1695379 | MI:0027 - cosedimentation |
| P26955 | P04401 | 8258700 | MI:0004 - affinity chromatography technologies |
| P26955 | P07321 | 1695379 | MI:0027 - cosedimentation |
| P26955 | P15247 | 1695379 | MI:0027 - cosedimentation |
| P27601 | Q76MZ3 | 15525651 | MI:0059 - gst pull down |
| P27699 | Q8R4I4 | 14512522 | MI:0018 - two hybrid |
| P28651 | Q9ERP3 | 18157088 | MI:0018 - two hybrid |
| P29351 | Q8C180 | 9632781 | MI:0019 - coimmunoprecipitation |
| P30658 | A2AM29 | 11439343 | MI:0018 - two hybrid |
| P30658 | P30658 | 9858600 | MI:0018 - two hybrid |
| P31786 | O89090 | 14530276 | MI:0040 - electron microscopy |
| P35279 | Q91V27 | 11856727 | MI:0007 - anti tag coimmunoprecipitation |
| P35285 | Q80T23 | 12051743 | MI:0096 - pull down |
| P35285 | Q91V27 | 11856727 | MI:0007 - anti tag coimmunoprecipitation |
| P35293 | Q80T23 | 12051743 | MI:0096 - pull down |
| P35293 | Q91V27 | 11856727 | MI:0007 - anti tag coimmunoprecipitation |
| P35295 | Q80T23 | 12051743 | MI:0096 - pull down |
| P35295 | Q91V27 | 11856727 | MI:0007 - anti tag coimmunoprecipitation |
| P35396 | Q6PDY0 | 15644333 | MI:0007 - anti tag coimmunoprecipitation |
| P35917 | Q00731 | 7970715 | MI:0004 - affinity chromatography technologies |
| P37889 | Q9QZJ6 | 17324935 | MI:0892 - solid phase assay |
| P39447 | Q8VHG2 | 16043488 | MI:0019 - coimmunoprecipitation |
| P41593 | Q9WTJ5 | 15823033 | MI:0019 - coimmunoprecipitation, MI:0113 - western blot |
| P42586 | Q62414 | 14573534 | MI:0019 - coimmunoprecipitation |
| P43407 | P21981 | 20929862 | MI:0019 - coimmunoprecipitation |
| P46414 | Q03147 | 7954814 | MI:0019 - coimmunoprecipitation |
| P46662 | Q61391 | 19388049 | MI:0019 - coimmunoprecipitation |
| P47708 | P61027 | 11773082 | MI:0007 - anti tag coimmunoprecipitation |
| P51141 | Q9WV60 | 10428961 | MI:0007 - anti tag coimmunoprecipitation |
| P51150 | Q80T23 | 12051743 | MI:0096 - pull down |
| P51150 | Q91V27 | 11856727 | MI:0007 - anti tag coimmunoprecipitation |
| P51680 | P10855 | 9192769 | MI:0405 - competition binding |
| P51680 | P14097 | 9192769 | MI:0405 - competition binding |
| P51680 | P30882 | 9192769 | MI:0405 - competition binding |
| P53566 | Q6PDY0 | 15644333 | MI:0007 - anti tag coimmunoprecipitation |
| P54130 | P16092 | 8619928 | MI:0004 - affinity chromatography technologies |
| P54130 | Q03142 | 8619928 | MI:0004 - affinity chromatography technologies |
| P55258 | Q80T23 | 12051743 | MI:0096 - pull down |
| P55258 | Q91V27 | 11856727 | MI:0007 - anti tag coimmunoprecipitation |
| P55258 | Q99N48 | 11773082 | MI:0007 - anti tag coimmunoprecipitation |
| P55258 | Q99N50 | 11773082 | MI:0007 - anti tag coimmunoprecipitation |
| P55258 | Q9R0Q1 | 11773082 | MI:0007 - anti tag coimmunoprecipitation |
| P56371 | Q80T23 | 12051743 | MI:0096 - pull down |
| P56371 | Q91V27 | 11856727 | MI:0007 - anti tag coimmunoprecipitation |
| P56371 | Q9D0C1 | 12972561 | MI:0096 - pull down; MI:0019 |
| P56546 | P30658 | 9858600 | MI:0018 - two hybrid |
| P61027 | Q80T23 | 12051743 | MI:0096 - pull down |
| P61027 | Q91V27 | 11856727 | MI:0007 - anti tag coimmunoprecipitation |
| P61027 | Q99N48 | 11773082 | MI:0007 - anti tag coimmunoprecipitation |
| P61027 | Q99N50 | 11773082 | MI:0007 - anti tag coimmunoprecipitation |
| P61027 | Q9R0Q1 | 11773082 | MI:0007 - anti tag coimmunoprecipitation |
| P61588 | P70336 | 16472685 | MI:0217 - phosphorylation reaction |
| P62259 | Q61097 | 10891492 | MI:0019 - coimmunoprecipitation |
| P62492 | Q80T23 | 12051743 | MI:0096 - pull down |
| P62492 | Q91V27 | 11856727 | MI:0007 - anti tag coimmunoprecipitation |

| | | | |
|---|---|---|---|
| P62500 | Q00322 | 15644333 | MI:0007 - anti tag coimmunoprecipitation |
| P62821 | Q80T23 | 12051743 | MI:0096 - pull down |
| P62821 | Q91V27 | 11856727 | MI:0007 - anti tag coimmunoprecipitation |
| P62880 | P63213 | 1629181 | MI:0004 - affinity chromatography technologies |
| P63011 | Q91V27 | 11856727 | MI:0007 - anti tag coimmunoprecipitation |
| P63011 | Q99N48 | 11773082 | MI:0007 - anti tag coimmunoprecipitation |
| P63011 | Q99N50 | 11773082 | MI:0007 - anti tag coimmunoprecipitation |
| P63011 | Q9R0Q1 | 11773082 | MI:0007 - anti tag coimmunoprecipitation |
| P63101 | Q61097 | 10891492 | MI:0019 - coimmunoprecipitation |
| P63239 | Q58A65 | 17074887 | MI:0018 - two hybrid |
| P63330 | Q9EP53 | 18291711 | MI:0019 - coimmunoprecipitation |
| P70196 | Q62523 | 17092936 | MI:0007 - anti tag coimmunoprecipitation; MI:0096 |
| P70196 | Q8BFW7 | 17092936 | MI:0007 - anti tag coimmunoprecipitation; MI:0096 |
| P70217 | P97471 | 16087734 | MI:0018 - two hybrid |
| P70217 | Q60954 | 15617687 | MI:0018 - two hybrid |
| P70371 | Q3UES3 | 12080061 | MI:0004 - affinity chromatography technologies |
| P70371 | Q6PFX9 | 12080061 | MI:0004 - affinity chromatography technologies |
| P70452 | Q549X6 | 16186111 | MI:0007 - anti tag coimmunoprecipitation |
| P97428 | Q9Z0G0 | 9770488 | MI:0018 - two hybrid |
| P97503 | Q8C2B3 | 14612411 | MI:0006 - anti bait coimmunoprecipitation |
| P97503 | Q9WU42 | 14612411 | MI:0006 - anti bait coimmunoprecipitation |
| Q01279 | Q9QUQ5 | 16144838 | MI:0006 - anti bait coimmunoprecipitation |
| Q02248 | Q02257 | 17925400 | MI:0006 - anti bait coimmunoprecipitation |
| Q02248 | Q8BN78 | 10207085 | MI:0019 - coimmunoprecipitation |
| Q04863 | Q04207 | 9070378 | MI:0004 - affinity chromatography technologies |
| Q08879 | Q9QZJ6 | 17324935 | MI:0892 - solid phase assay |
| Q09163 | Q01705 | 20457810 | MI:0019 - coimmunoprecipitation |
| Q505F1 | Q60953 | 19204783 | MI:0019 - coimmunoprecipitation |
| Q548T0 | Q549X6 | 16186111 | MI:0007 - anti tag coimmunoprecipitation |
| Q548T0 | Q80T23 | 16186111 | MI:0007 - anti tag coimmunoprecipitation |
| Q548T0 | Q99N48 | 16186111 | MI:0007 - anti tag coimmunoprecipitation |
| Q548T0 | Q99N50 | 16186111 | MI:0007 - anti tag coimmunoprecipitation |
| Q549X6 | Q60770 | 16186111 | MI:0007 - anti tag coimmunoprecipitation |
| Q60770 | Q8K1E0 | 16186111 | MI:0007 - anti tag coimmunoprecipitation |
| Q60972 | Q01147 | 10866654 | MI:0018 - two hybrid |
| Q61146 | Q8VHG2 | 16043488 | MI:0019 - coimmunoprecipitation |
| Q61205 | Q924X6 | 17330141 | MI:0019 - coimmunoprecipitation |
| Q61206 | Q924X6 | 17330141 | MI:0019 - coimmunoprecipitation |
| Q61559 | P27512 | 21148035 | MI:0110 - text mining, MI:0114 x-ray crystallography |
| Q61730 | P01582 | 17669273 | MI:0030 - cross-linking study |
| Q62009 | Q08024 | 9632804 | MI:0004 - affinity chromatography technologies |
| Q62108 | P28652 | 19455133 | MI:0676 - tandem affinity purification |
| Q62312 | Q8CIN4 | 14612425 | MI:0006 - anti bait coimmunoprecipitation |
| Q62424 | P97471 | 16087734 | MI:0018 - two hybrid |
| Q63912 | Q7M6Z0 | 17189258 | MI:0415 - enzymatic studies |
| Q63912 | Q8K0S5 | 17189258 | MI:0415 - enzymatic studies |
| Q64092 | Q03141 | 20214879 | MI:0019 - coimmunoprecipitation |
| Q64343 | Q9DBM0 | 14504269 | MI:0019 - coimmunoprecipitation |
| Q64729 | Q8CIN4 | 14612425 | MI:0006 - anti bait coimmunoprecipitation |
| Q66T02 | P60766 | 20811643 | MI:0096 - pull down |
| Q6NXK8 | Q62108 | 19571134 | MI:0019 - coimmunoprecipitation |
| Q6TDP3 | Q9WTX6 | 17062563 | MI:0019 - coimmunoprecipitation |
| Q6Y5D8 | Q8CIN4 | 115471851 | MI:0018 - two hybrid |
| Q71VB4 | Q7TPV4 | 17875935 | MI:0019 - coimmunoprecipitation |
| Q7M6Z0 | Q8K0T0 | 17189258 | MI:0415 - enzymatic studies |
| Q7M6Z0 | Q99P72 | 17189258 | MI:0415 - enzymatic studies |
| Q7M6Z0 | Q9ES97 | 17189258 | MI:0415 - enzymatic studies |
| Q7TMS5 | Q9DBM0 | 14504269 | MI:0019 - coimmunoprecipitation |
| Q80T23 | Q99KL7 | 12051743 | MI:0096 - pull down |
| Q80T23 | Q9CQD1 | 12051743 | MI:0096 - pull down |
| Q80T23 | Q9R0M6 | 12051743 | MI:0096 - pull down |
| Q80T23 | Q9WTL2 | 12051743 | MI:0096 - pull down |
| Q8BGS1 | P62331 | 18794329 | MI:1016 - fluorescence recovery after photobleaching |
| Q8BPB5 | Q9QZJ6 | 17324935 | MI:0892 - solid phase assay |
| Q8CBD1 | P68254 | 18628823 | MI:0019 - coimmunoprecipitation |
| Q8CBD1 | Q9JIF0 | 18628823 | MI:0019 - coimmunoprecipitation, MI:0110 - text mining |
| Q8CF89 | Q62312 | 19556242 | MI:0019 - coimmunoprecipitation |
| Q8CF89 | Q64729 | 19556242 | MI:0019 - coimmunoprecipitation |
| Q8CIN4 | Q8CIN4 | 115471851 | MI:0018 - two hybrid |
| Q8K0S5 | Q8K0T0 | 17189258 | MI:0415 - enzymatic studies |
| Q8K0S5 | Q99P72 | 17189258 | MI:0415 - enzymatic studies |

| | | | |
|---|---|---|---|
| Q8K0S5 | Q9ES97 | 17189258 | MI:0415 - enzymatic studies |
| Q8K0T0 | Q99PI8 | 17189258 | MI:0415 - enzymatic studies |
| Q8R4G0 | O08747 | 14595443 | MI:0004 - affinity chromatography technologies |
| Q8R4G0 | Q8K1S3 | 14595443 | MI:0004 - affinity chromatography technologies |
| Q8R4G0 | Q8K1S4 | 14595443 | MI:0004 - affinity chromatography technologies |
| Q8VHZ5 | Q9EP53 | 19129461 | MI:0019 - coimmunoprecipitation |
| Q91V27 | Q99KL7 | 11856727 | MI:0007 - anti tag coimmunoprecipitation |
| Q91V27 | Q9CQD1 | 11856727 | MI:0007 - anti tag coimmunoprecipitation |
| Q91V27 | Q9R0M6 | 11856727 | MI:0007 - anti tag coimmunoprecipitation |
| Q91V27 | Q9WTL2 | 11856727 | MI:0007 - anti tag coimmunoprecipitation |
| Q922H7 | O35182 | 15239668 | MI:0019 - coimmunoprecipitation |
| Q922H7 | O35253 | 15239668 | MI:0019 - coimmunoprecipitation |
| Q922H7 | P97471 | 15239668 | MI:0019 - coimmunoprecipitation |
| Q922H7 | Q62432 | 15239668 | MI:0019 - coimmunoprecipitation |
| Q922H7 | Q8BUN5 | 15239668 | MI:0019 - coimmunoprecipitation |
| Q99K43 | P38532 | 18570919 | MI:0018 - two hybrid |
| Q9EQW6 | Q99MA9 | 14573534 | MI:0019 - coimmunoprecipitation |
| Q9ERI2 | Q99104 | 12006666 | MI:0004 - affinity chromatography technologies |
| Q9ERP3 | Q9JK37 | 18157088 | MI:0018 - two hybrid |
| Q9JM90 | Q9QVP9 | 10679268 | MI:0018 - two hybrid |
| Q9JMB0 | P05132 | 10671526 | MI:0018 - two hybrid |
| Q9JMB0 | P0C605 | 10671526 | MI:0018 - two hybrid |
| Q9QXX8 | Q61584 | 10556305 | MI:0018 - two-hybrid, MI:0059 - gst pull down |
| Q9QZJ6 | Q9WVH9 | 17324935 | MI:0892 - solid phase assay |
| Q9QZJ6 | Q9WVJ9 | 17324935 | MI:0892 - solid phase assay |
| Q9WTW2 | P23299 | 19623261 | MI:0019 - coimmunoprecipitation |
| O08599 | P70452 | 16186111 | MI:0007 - anti tag coimmunoprecipitation |

**Table S3. CORUM database complexes from mouse (Mus musculus).**

| **Parvulin-associated pre-rRNP**<br>complex ID: 3047<br>PubMed ID: 11960984 | | **Respiratory chain complex I**<br>complex ID: 382<br>PubMed ID: 15591592 | | **20S Immuno-proteasome**<br>complex ID: 39<br>PubMed ID: 10436176 | | **c-Src immune complex**<br>complex ID: 6938<br>PubMed ID: 19628583 | | **DNA synthesome complex**<br>complex ID: 1001<br>PubMed ID: 8126085 | |
|---|---|---|---|---|---|---|---|---|---|
| **Protein** | **Name** | **Protein** | **Name** | **Protein** | **Name** | **Protein** | **Name** | **Protein** | **Name** |
| O09167 | Rpl21 | O09111 | Ndufb11 | O09061 | Psmb1 | P05213 | Tuba1b | O35654 | Pold2 |
| O43143 | DHX15 | O35683 | Ndufa1 | O35955 | Psmb10 | P05480 | Src | P17918 | Pcna |
| O76021 | RSL1D1 | P03888 | Mtnd1 | O70435 | Psma3 | P16054 | Prkce | P20664 | Prim1 |
| O95995 | GAS8 | P03899 | Mtnd3 | P28063 | Psmb8 | P20444 | Prkca | P33609 | Pola1 |
| P09405 | Ncl | P03911 | Mtnd4 | P28076 | Psmb9 | P28867 | Prkcd | P33610 | Prim2 |
| P11276 | Fn1 | P52503 | Ndufs6 | P49722 | Psma2 | P34152 | Ptk2 | P33611 | Pola2 |
| P12970 | Rpl7a | Q3UIU2 | Ndufb6 | P99026 | Psmb4 | P49817 | Cav1 | P37913 | Lig1 |
| P14148 | Rpl7 | Q62425 | Ndufa4 | Q9QUM9 | Psma6 | P63085 | Mapk1 | P52431 | Pold1 |
| P14869 | Rplp0 | Q7TMF3 | Ndufa12 | Q9R1P0 | Psma4 | P68369 | Tuba1a | Q01320 | Top2a |
| P19253 | Rpl13a | Q8K3J1 | Ndufs8 | Q9R1P1 | Psmb3 | Q01776 | Gnrhr | Q04750 | Top1 |
| P21127 | CDK11B | Q91VD9 | Ndufs1 | Q9R1P3 | Psmb2 | Q61097 | Ksr1 | Q9CWP8 | Pold4 |
| P27635 | RPL10 | Q91WD5 | Ndufs2 | Q9R1P4 | Psma1 | Q63844 | Mapk3 | Q9EQ28 | Pold3 |
| P27659 | Rpl3 | Q91WP8 | Ndufv3 | Q9Z2U0 | Psma7 | Q64727 | Vcl | | |
| P35550 | Fbl | Q91YT0 | Ndufv1 | Q9Z2U1 | Psma5 | Q8VI36 | Pxn | | |
| P35980 | Rpl18 | Q99LC3 | Ndufa10 | | | | | | |
| P46781 | RPS9 | Q99LY9 | Ndufs5 | | | | | | |
| P47911 | Rpl6 | Q9CPP6 | Ndufa5 | | | | | | |
| P47962 | Rpl5 | Q9CPU2 | Ndufb2 | | | | | | |
| P47963 | Rpl13 | Q9CQ54 | Ndufc2 | | | | | | |
| P53569 | Cebpz | Q9CQ75 | Ndufa2 | | | | | | |
| P56183 | Rrp1 | Q9CQ91 | Ndufa3 | | | | | | |
| P61255 | Rpl26 | Q9CQC7 | Ndufb4 | | | | | | |
| P62242 | Rps8 | Q9CQH3 | Ndufb5 | | | | | | |
| P62702 | Rps4x | Q9CQJ8 | Ndufb9 | | | | | | |
| P62717 | Rpl18a | Q9CQZ5 | Ndufa6 | | | | | | |
| P62754 | Rps6 | Q9CQZ6 | Ndufb3 | | | | | | |
| P62906 | RPL10A | Q9CR21 | Ndufab1 | | | | | | |
| P62908 | Rps3 | Q9CR61 | Ndufb7 | | | | | | |
| P62918 | Rpl8 | Q9CXZ1 | Ndufs4 | | | | | | |
| P68363 | TUBA1B | Q9D6J5 | Ndufb8 | | | | | | |
| P84099 | Rpl19 | Q9D6J6 | Ndufv2 | | | | | | |
| P97351 | Rps3a | Q9D8B4 | Ndufa11 | | | | | | |
| P97452 | Bop1 | Q9DC69 | Ndufa9 | | | | | | |
| P99024 | Tubb5 | Q9DC70 | Ndufs7 | | | | | | |
| Q61656 | Ddx5 | Q9DCJ5 | Ndufa8 | | | | | | |
| Q6DFW4 | Nop58 | Q9DCS9 | Ndufb10 | | | | | | |
| Q7TPV4 | Mybbp1a | Q9DCT2 | Ndufs3 | | | | | | |
| Q8K363 | Ddx18 | Q9ERS2 | Ndufa13 | | | | | | |
| Q91VE6 | Nifk | Q9MD82 | mt-Nd5 | | | | | | |
| Q921N6 | Ddx27 | Q9Z1P6 | Ndufa7 | | | | | | |
| Q922K7 | Nop2 | | | | | | | | |
| Q99LH1 | Gnl2 | | | | | | | | |
| Q99ME9 | Gtpbp4 | | | | | | | | |
| Q9BWT6 | MND1 | | | | | | | | |
| Q9CR57 | Rpl14 | | | | | | | | |
| Q9CYH6 | Rrs1 | | | | | | | | |
| Q9CZM2 | Rpl15 | | | | | | | | |
| Q9D0I8 | Mrto4 | | | | | | | | |
| Q9D0R4 | Ddx56 | | | | | | | | |
| Q9D6Z1 | Nop56 | | | | | | | | |
| Q9D8E6 | Rpl4 | | | | | | | | |
| Q9D8N0 | Eef1g | | | | | | | | |
| Q9D903 | Ebna1bp2 | | | | | | | | |
| Q9DBE9 | Ftsj3 | | | | | | | | |
| Q9EQ61 | Pes1 | | | | | | | | |
| Q9ESV0 | Ddx24 | | | | | | | | |
| Q9JIK5 | Ddx21 | | | | | | | | |
| Q9JJ80 | Rpf2 | | | | | | | | |
| Q9JJA4 | Wdr12 | | | | | | | | |
| Q9NW13 | RBM28 | | | | | | | | |
| Q9ULW0 | TPX2 | | | | | | | | |
| Q9Y237 | PIN4 | | | | | | | | |

**Table S4. Proteins identified to be localized to lipid rafts upon B cell activation.**

| Proteins 1-65 | | Proteins 66-130 | | Proteins 131-195 | | Proteins196-260 | | Proteins 261-324 | |
|---|---|---|---|---|---|---|---|---|---|
| **UniProt** | **Name** | **UniProt** | **Name** | **UniProt** | **Name** | **UniProt** | **Name** | **UniProt** | **Name** |
| Q8BGQ7 | Aars | Q9WVK4 | Ehd1 | Q8R081 | Hnrnpl | Q9ERS5 | Plekha2 | P11157 | Rrm2 |
| Q99LE6 | Abcf2 | Q8JZQ9 | Eif3b | P07901 | Hsp90aa1 | Q9D3P8 | Plgrkt | Q99LF4 | Rtcb |
| Q8K268 | Abcf3 | Q8R1B4 | Eif3c | P20029 | Hspa5 | Q923G2 | Polr2h | Q99P72 | Rtn4 |
| Q91V92 | Acly | P60229 | Eif3e | P63017 | Hspa8 | Q91YU8 | Ppan | P60122 | Ruvbl1 |
| Q9D358 | Acp1 | Q8QZY1 | Eif3l | Q61699 | Hsph1 | P62137 | Ppp1ca | Q3UU41 | Scimp |
| Q9QUJ7 | Acsl4 | Q99JX4 | Eif3m | Q6ZQA6 | Igsf3 | P63330 | Ppp2ca | O08547 | Sec22b |
| P60710 | Actb | P60843 | Eif4a1 | O54824 | Il16 | P35700 | Prdx1 | Q9CY58 | Serbp1 |
| P63260 | Actg1 | Q8BGD9 | Eif4b | P24547 | Impdh2 | Q61171 | Prdx2 | Q8VIJ6 | Sfpq |
| P57780 | Actn4 | Q6NZJ6 | Eif4g1 | Q9EPL8 | Ipo7 | O08807 | Prdx4 | Q9ET39 | Slamf6 |
| P61161 | Actr2 | Q80XI3 | Eif4g3 | Q9JKF1 | Iqgap1 | P68404 | Prkcb | P53986 | Slc16a1 |
| Q8BK64 | Ahsa1 | P63242 | Eif5a | P81122 | Irs2 | Q8CIG8 | Prmt5 | Q8JZU2 | Slc25a1 |
| O08915 | Aip | P70372 | Elavl1 | Q9JHU9 | Isyna1 | Q9D7G0 | Prps1 | Q8BH59 | Slc25a12 |
| P05064 | Aldoa | Q8BPU7 | Elmo1 | O54890 | Itgb3 | Q8BHE0 | Prr11 | Q9Z2Z6 | Slc25a20 |
| P97384 | Anxa11 | P21995 | Emb | Q3U0V1 | Khsrp | P62192 | Psmc1 | Q8VEM8 | Slc25a3 |
| Q07076 | Anxa7 | P17182 | Eno1b | Q61768 | Kif5b | O88685 | Psmc3 | P48962 | Slc25a4 |
| P84078 | Arf1 | P48193 | Epb4.1 | P52293 | Kpna2 | P62196 | Psmc5 | P10852 | Slc3a2 |
| P61750 | Arf4 | O70318 | Epb4.1l2 | P70168 | Kpnb1 | P62334 | Psmc6 | Q99JR8 | Smarcd2 |
| P62331 | Arf6 | Q8BGS1 | Epb4.1l5 | Q61595 | Ktn1 | Q8BG32 | Psmd11 | Q8CG48 | Smc2 |
| Q3UIA2 | Arhgap17 | Q8CGC7 | Eprs | Q8BMJ2 | Lars | Q99JI4 | Psmd6 | Q9CW03 | Smc3 |
| Q8VDN2 | Atp1a1 | P42567 | Eps15 | Q3U9G9 | Lbr | P29351 | Ptpn6 | P62320 | Snrpd3 |
| O55143 | Atp2a2 | Q61545 | Ewsr1 | Q61233 | Lcp1 | O35295 | Purb | Q3UMC0 | Spata5 |
| Q9Z1G3 | Atp6v1c1 | P26040 | Ezr | Q99N69 | Lpxn | Q8BML9 | Qars | Q9CYN2 | Spcs2 |
| P28658 | Atxn10 | Q6A0A9 | Fam120a | P19973 | Lsp1 | Q91V41 | Rab14 | Q64337 | Sqstm1 |
| Q7TQH0 | Atxn2l | P19096 | Fasn | Q01965 | Ly9 | P35293 | Rab18 | Q6P069 | Sri |
| B2RQC6 | Cad | P35550 | Fbl | Q8C052 | Map1s | P62821 | Rab1 | Q60864 | Stip1 |
| P35564 | Canx | P08101 | Fcgr2b | Q68FL6 | Mars | Q9D1G1 | Rab1b | P46978 | Stt3a |
| P47754 | Capza2 | Q8K1B8 | Fermt3 | Q8VCF0 | Mavs | P35282 | Rab21 | Q6A028 | Swap70 |
| P47757 | Capzb | Q69ZL1 | Fgd6 | Q9JKP5 | Mbnl1 | Q9CQD1 | Rab5a | Q7TMK9 | Syncrip |
| Q9WU84 | Ccs | P30416 | Fkbp4 | P25206 | Mcm3 | P35278 | Rab5c | Q9WVA4 | Tagln2 |
| P80315 | Cct4 | Q8BTM8 | Flna | P49717 | Mcm4 | P35279 | Rab6a | Q921F2 | Tardbp |
| P42932 | Cct8 | Q80X90 | Flnb | Q61881 | Mcm7 | P51150 | Rab7 | P10711 | Tcea1 |
| P25918 | Cd19 | P09528 | Fth1 | Q9CQT1 | Mri1 | P63001 | Rac1 | Q62351 | Tfrc |
| Q61470 | Cd37 | Q91WJ8 | Fubp1 | P19437 | Ms4a1 | Q05144 | Rac2 | Q9D880 | Timm50 |
| P21855 | Cd72 | Q91Z49 | Fyttd1 | P26041 | Msn | Q9WVM1 | Racgap1 | P40142 | Tkt |
| P40237 | Cd82 | P97379 | G3bp2 | P00405 | mt-Co2 | Q9ERU9 | Ranbp2 | P26039 | Tln1 |
| P60766 | Cdc42 | Q9CZD3 | Gars | Q922D8 | Mthfd1 | Q8BMS9 | Rassf2 | Q61029 | Tmpo |
| P11440 | Cdk1 | Q64737 | Gart | Q8R1S4 | Mtss1 | Q91VM5 | Rbmxl1 | Q8BFY9 | Tnpo1 |
| Q4VAA2 | Cdv3 | Q8VHR5 | Gatad2b | Q7TPV4 | Mybbp1a | Q8BK67 | Rcc2 | P17751 | Tpi1 |
| Q8BT07 | Cep55 | O09172 | Gclm | Q99KQ4 | Nampt | Q9WUK4 | Rfc2 | P21107 | Tpm3 |
| P18760 | Cfl1 | E9PVA8 | Gcn1l1 | Q8C156 | Ncaph | Q6A0D4 | Rftn1 | Q64514 | Tpp2 |
| Q9CRB9 | Chchd3 | Q61598 | Gdi2 | P46935 | Nedd4 | Q9QUI0 | Rhoa | P63028 | Tpt1 |
| Q9Z1Q5 | Clic1 | Q6Y7W8 | Gigyf2 | Q8K4Q6 | Neil1 | P84096 | Rhog | P99024 | Tubb5 |
| Q9QYB1 | Clic4 | Q9WU65 | Gk2 | Q9WTK5 | Nfkb2 | Q9QZL0 | Ripk3 | P10639 | Txn1 |
| Q68FD5 | Cltc | Q3THK7 | Gmps | Q01768 | Nme2 | Q6ZWV3 | Rpl10 | Q9D883 | U2af1 |
| Q9DBP5 | Cmpk1 | P27601 | Gna13 | Q8K2T1 | Nmral1 | P84099 | Rpl19 | P0DP28 | unknown |
| P53996 | Cnbp | P08752 | Gnai2 | Q61937 | Npm1 | P62830 | Rpl23 | Q02053 | Uba1 |
| P16330 | Cnp | Q9DC51 | Gnai3 | Q9R1J0 | Nsdhl | P27659 | Rpl3 | Q91VX2 | Ubap2 |
| O88207 | Col5a1 | P63094 | Gnas | Q5F2E7 | Nufip2 | P62889 | Rpl30 | Q80X50 | Ubap2l |
| O89053 | Coro1a | P62880 | Gnb2 | Q80U93 | Nup214 | Q9D8E6 | Rpl4 | Q9EPU0 | Upf1 |
| Q8CJ40 | Crocc | Q9QYE6 | Golga5 | Q8BJ71 | Nup93 | P47911 | Rpl6 | P52479 | Usp10 |
| P70698 | Ctps | Q8BUV3 | Gphn | Q7TQI3 | Otub1 | P14148 | Rpl7 | Q3UJD6 | Usp19 |
| Q922B2 | Dars | O70325 | Gpx4 | B2RRE7 | Otud4 | Q91YQ5 | Rpn1 | Q9Z1Q9 | Vars |
| Q91VR5 | Ddx1 | Q99L20 | Gstt3 | P29341 | Pabpc1 | P63325 | Rps10 | Q01853 | Vcp |
| Q501J6 | Ddx17 | P53702 | Hccs | Q9DCL9 | Paics | P62245 | Rps15a | Q60930 | Vdac2 |
| Q62167 | Ddx3x | P49710 | Hcls1 | Q3TC46 | Patl1 | P14131 | Rps16 | Q60931 | Vdac3 |
| P54823 | Ddx6 | Q9JKY5 | Hip1r | P60335 | Pcbp1 | P63276 | Rps17 | P62960 | Ybx1 |
| P00375 | Dhfr | Q00547 | Hmmr | Q9WU78 | Pdcd6ip | P60867 | Rps20 | P59326 | Ythdf1 |
| Q811D0 | Dlg1 | O70252 | Hmox2 | Q922R8 | Pdia6 | P62849 | Rps24 | Q91YT7 | Ythdf2 |
| P63037 | Dnaja1 | P49312 | Hnrnpa1 | Q8R1G6 | Pdlim2 | Q6ZWU9 | Rps27rt | Q9CQV8 | Ywhab |
| Q9QYJ0 | Dnaja2 | O88569 | Hnrnpa2b1 | Q9WUA3 | Pfkp | Q6ZWY3 | Rps27l | P62259 | Ywhae |
| Q8C147 | Dock8 | Q8BG05 | Hnrnpa3 | P62962 | Pfn1 | P97461 | Rps5 | P68510 | Ywhah |
| Q9JHU4 | Dync1h1 | Q99020 | Hnrnpab | Q9JJV2 | Pfn2 | P62754 | Rps6 | P63101 | Ywhaz |
| P57776 | Eef1d | Q9Z2X1 | Hnrnpf | Q61753 | Phgdh | P14206 | Rpsa | Q3UPF5 | Zc3hav1 |
| Q9D8N0 | Eef1g | O35737 | Hnrnph1 | Q7M6Y3 | Picalm | P62071 | Rras2 | B2RX14 | Zcchc11 |
| P58252 | Eef2 | P61979 | Hnrnpk | Q8CIH5 | Plcg2 | P07742 | Rrm1 | | |

**Table S5. Proteins predicted to co-evolve with Cad.**

| **UniProt ID** | **Name** | **Residues co-evolving with Cad** | **Subcellular location** |
|---|---|---|---|
| Q8JZQ9 | Eif3b | 207I | Cytoplasm |
| Q8CGC7 | Eprs | 5C, 12N, 747A | Cytoplasm, Membrane |
| P19096 | Fasn | 2268Q | Cytoplasm; GO: Golgi, Mitochondrion, membrane |
| P19437 | Ms4a1 | 191M, 213M, 219S | Membrane (multipass); GO: Nucleoplasm |
| Q80U93 | Nup214 | 1513A, 1511P | Nuclear pore (cytoplasmic side) |
| Q9ERU9 | Ranbp2 | 229R | Nucleus, Nuclear pore, Nuclear envelope; GO: mitochondrion |
| Q91YQ5 | Rpn1 | 89I, 86G, 381N, 389D, 491L | ER; GO: Cytosol, |
| Q99P72 | Rtn4 | 248T, 263R, 266N, 309V, 324H | Membrane, ER; GO: Nuclear envelope |
| P10852 | Slc3a2 | 165I | Membrane, Lysosome membrane; GO: nucleoplasm |
| Q3UMC0 | Spata5 | 507A, 538R, 589S, 609I | Mitochondrion, Cytoplasm, Spindle |
| **UniProt** | **Name** | **Residues co-evolving from Cad** | **Subcellular location** |
| B2RQC6 | **Cad** | 20A, 1728L, 1887G, 1892A, 2108S, 2114S, 2610A | Cytoplasm, Nucleus |
